# New insights underlying the early events of dopaminergic dysfunction in Parkinson’s Disease

**DOI:** 10.1101/2020.09.27.313957

**Authors:** Hannah L. Dela Cruz, Esther L. Dela Cruz, Cody J. Zurhellen, Herbert T. York, Jim A. Baun, Joshua L. Dela Cruz, Jay S. Dela Cruz

## Abstract

Alpha melanocyte-stimulating hormone (**α**-MSH) is an autocrine factor released by activated microglia during neuroinflammation and is elevated in the cerebrospinal fluid of Parkinson’s disease (PD) patients. **α**-MSH impaired cellular autophagy and induced the accumulation of alpha-synuclein in a melanized human dopaminergic cell model. Increased **α**-MSH in the brain of mice resulted in the gradual worsening of abnormal gait. Dopamine replacement with L-dopa/Benserazide or treatment with a dopamine receptor agonist, Pramipexole, temporarily restored normal gait, suggesting dopamine deficiency as the cause of motor deficits in these mice. Notably, end-stage disease pathology such as neuronal cell loss, reduction in tyrosine hydroxylase (TH)+ fiber density in the striatum and pSer129+ alpha-synuclein inclusions were absent. Rather, autophagic dysfunction was observed in the dopaminergic neuronal (DN) cell population within the substantia nigra pars compacta and ventral tegmental area. Moreover, increased expression of TH was observed in the striatum, suggesting a compensatory response to diminished dopamine levels. Our findings provide new insights into the early events that underlie neurodegeneration in PD and suggest that exposure of DNs to elevated levels of microglial **α**-MSH leads to impairment of autophagy resulting in abnormal accumulation of proteins, dopaminergic dysfunction and motor deficits.

**Graphical abstract:** 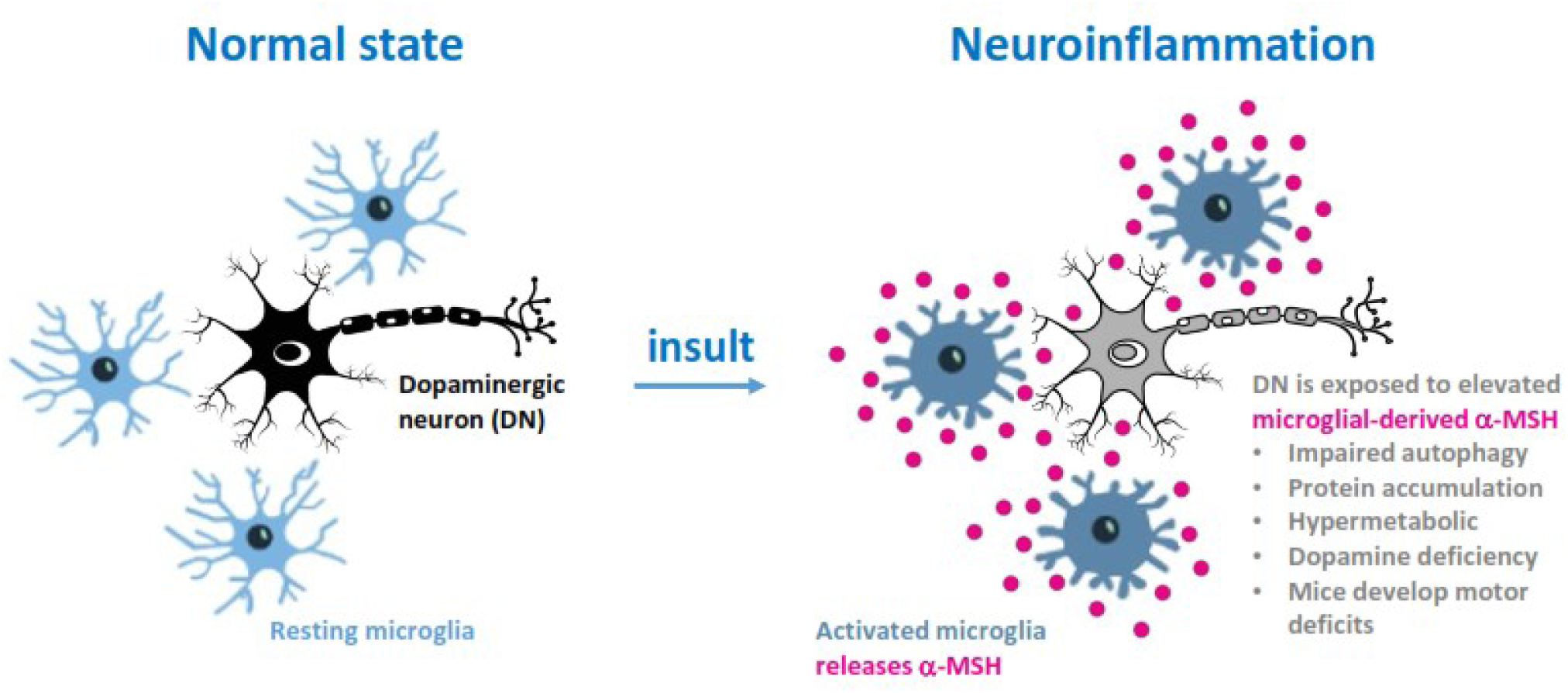

**Significance statement:** We now show that a naturally occurring compound increased in the brain of Parkinson’s disease (PD) patients, called **α**-MSH, can trigger abnormal accumulation of alpha-synuclein in a dopaminergic cell model. Increasing **α**-MSH in the brain of mice resulted in motor symptoms and abnormal gait. Increasing dopamine activity in these mice using Levodopa or Pramipexole restored normal gait, suggesting that the mice were deficient in dopamine, as seen in PD. We now describe a cell and an animal model that can reproduce the early stages of dopaminergic dysfunction in PD. These new pre-clinical research tools will be useful in developing effective drugs that will stop the progression of the disease in patients who suffer from PD.

**Abbreviations:** PD, Parkinson’s disease; DN, dopaminergic neuron; **α**-MSH, alpha-melanocyte stimulating hormone; TH, tyrosine hydroxylase; SNpc, substantia nigra pars compacta; VTA, ventral tegmental area; CNS, central nervous system; CSF, cerebrospinal fluid; INS, intranasal; ASIP, agouti-signaling protein; MC1R, melanocortin receptor 1; ROS, reactive-oxygen species; MSA, multiple system atrophy

## Introduction

The events that initiate and drive dopaminergic dysfunction in Parkinson’s disease (PD) remain unclear. A leading hypothesis suggests that pathologic misfolded alpha-synuclein is released by affected cells in the brain and the disease is spread to healthy cells in a prion-like manner [1]. While there is still no proof that this mechanism of transmission and propagation of pathology is active in the brains of PD patients, there is indirect evidence that suggests it is plausible [1]. Still, the more fundamental question of what triggers proteins to accumulate and misfold in dopaminergic neurons (DNs) in PD remains unanswered [2].

Abnormal protein accumulation suggests a failure with the cell’s autophagic machinery to recycle excess cellular components [3]. Protein accumulation is observed in DNs in PD as seen in aggregated cellular debris and proteins clumped in Lewy bodies and inclusions in post-mortem tissue [4]. Histology of PD substantia nigra (SN) showed autophagic degeneration, as seen in the accumulation of the autophagy flux marker, p62 protein, and reduction in markers of autophagy including LAMP-1 [5–7]. Disruption of autophagy in mice results in the formation of alpha-synuclein inclusions in DNs, neuronal cell loss and motor deficits [8, 9]. Consistent with these observations, mutated genes associated with familial PD (e.g., *SNCA*, *GBA* and *LRRK2*) have protein products that have important functions in cellular autophagy [10].

Neuroinflammation is also involved in PD pathophysiology and is mediated by activated microglia, which are the primary resident immune cells of the central nervous system (CNS) [11]. Activated microglia are present prior to development of alpha-synuclein inclusions in PD and neuronal cell loss *in-vivo* [12–14]. Indeed, injection of bacterial lipopolysaccharide in the SN of mice led to activation of microglia and subsequent formation of alpha-synuclein inclusions in DNs and eventual cell loss [15, 16]. While the importance microglial cells in neurodegenerative diseases is widely appreciated, the manner by which activated microglia drive neurodegeneration in PD remains unclear [13].

In spite of the complementary roles of neuroinflammation and dysregulation of autophagy in PD pathophysiology, the direct interplay between them is poorly understood [17]. In the course of our work to establish *in-vitro* cellular models more relevant to PD, we observed the 13 amino-acid peptide, alpha melanocyte-stimulating hormone (**α**-MSH), to impair cellular autophagy. **α**-MSH is an anorexigenic hormone expressed in the hypothalamic arcuate nucleus and in the nucleus tractus solitarius of the brainstem where it regulates energy homeostasis and cellular metabolism [18, 19]. It also functions as a potent anti-inflammatory mediator released by activated microglial cells to maintain homeostasis in the brain during neuroinflammation [20–22].

Treatment of PD patients with beta-MSH (similar in function to **α**-MSH) aggravated clinical symptoms [23], presumably by reducing striatal dopamine levels [24, 25]. In line with this observation, pharmacologic reduction of **α**-MSH resulted in improvements in PD symptoms [26–28]. Furthermore, **α**-MSH is elevated in the cerebrospinal fluid (CSF) of patients with PD [29, 30]. On the basis of these observations and our findings, we propose that **α**-MSH provides a fundamental link between neuroinflammation and dysregulation of autophagy in PD pathophysiology.

Below we describe our investigation into the dysregulation of autophagy by **α**-MSH using the melanized dopaminergic cell model, MNT-1. We observed several features of PD-associated pathology and metabolic features after exposure of MNT-1 to **α**-MSH, including impairment of autophagy, loss of pigmentation, accumulation of alpha-synuclein and hypermetabolism. Moreover, prolonged central exposure of normal mice to **α**-MSH resulted in a time-dependent worsening of motor deficits that was temporarily alleviated with dopamine replacement or treatment with a dopamine receptor agonist, suggesting the induction of dopaminergic dysfunction in these mice.

Histopathology analysis showed neither nigral cell loss, nor a decrease in striatal TH density, nor the appearance of alpha-synuclein inclusions in the substantia nigra. Instead, we observed autophagic impairment within the DNs in the substantia nigra pars compacta (SNpc)-VTA region and a compensatory response to low dopamine levels in the striatum of affected mice. These observations suggest that exposure of healthy DNs to elevated levels of **α**-MSH is sufficient to initiate dopaminergic dysfunction, which we propose underlies the early stages of neurodegeneration in PD.

## Materials and Methods

### Growth/culture conditions

MNT-1 was a gift from Dr. Michael D. Marks (University of Pennsylvania Perelman School of Medicine). MNT-1 growth media reagents were purchased from Thermofisher.com: DMEM (11995-065) supplemented with 15% FBS (10437-028), 10% MEM NEAA (1037-021), 10% AIM V (31035-025), and penicillin/streptomycin (from 100X, 15140-122). Cells were maintained in 25 cm^2^ canted neck tissue culture flask (353109) in 4 ml of media in a 37°C humidified incubator and 5% CO_2_. Passaging was done once confluency reached 90% by trypsinization (0.25% Trypsin, 15050-057) and neutralization in growth media. Cells were diluted 1:3 or 1:6, depending on need, and seeded in 4ml growth media. Growth media was replaced every 2-3 days. For cell assays, confluency of 80-90% was reach before cells were trypsinized and washed once in growth media and resuspended in the appropriate assay buffer. Cell counts were done in trypan blue using a hemacytometer under a microscope. **α**-MSH (GenScript, RP10644) was prepared in artificial cerebrospinal fluid (aCSF, Tocris, 3525) followed by removal of endotoxin (<0.01EU/ml, Pierce, 88274). Recombinant human (rh)ASIP (R&D Systems, 9094-AG) was prepared in aCSF.

### Visual observation of reduced melanin formation in **α**-MSH-treated MNT-1 cells

20,000 MNT-1 cells were seeded per well in a 24-well flat bottom tissue culture plate (Costar 353047) in 500 microliters of growth media and the plate placed in a 37°C humidified incubator with 5% CO_2_. After 48 hours, 250 microliters of supernatant was removed and replaced with the same volume of growth media or growth media containing 200nM **α**-MSH (final=100nM). After 9 days, the plate was placed under ambient light and a photograph taken using a digital camera.

### Measurement of melanin in a 96-well plate format

A) Conventional method: Supernatant from wells containing adherent MNT-1 was removed and replaced with 200 microliters of 1X PBS. 1X PBS was removed and the plate dabbed upside-down and allowed to sit this way over a stack of paper towels for 2 minutes. After another dab, 100 microliters of 1N NaOH was added to each well and the plate moved to a plate shaker (200 rpm, 5 minutes). The plate was moved to 37°C incubator (16-18 hours). Samples were further solubilized by resuspension and the absorbance measured at 405nm using a plate reader (Molecular Devices Thermomax and Softmax Pro software). The absorbance values from triplicate wells were averaged and the mean and standard deviation reported. B) Alternative method using fluorescence (fluorescein) quenching by melanin: Supernatant from wells containing adherent MNT-1 was removed and replaced with 200 microliters of 1X PBS. 1X PBS was removed and the plate dabbed upside-down and allowed to sit this way over a stack of paper towels for 2 minutes. The plate was frozen overnight in below −70°C and thawed at ambient room temperature for 20 minutes. A fluorescein-based dye was prepared in a cell lysis solution and 200 microliters added to each well and the plate placed on a shaker in the dark for 10 minutes. The lysed cells were resuspended and 90 microliters transferred to a white 96-well plate (Costar 353296) and fluorescence measured using a fluorescence plate reader (Cytofluor 4000, Gain 50, Ex 485/20, Em 530/25). The fluorescence signal is inversely proportional to the amount of melanin present in the well. The higher the signal the lower the amount of melanin that is present in the well. To obtain a reportable value, the fluorescence signal from the control wells (cells in medium alone) was divided by the fluorescence signal from test wells. The ratios (%) obtained from triplicate determinations were averaged and the mean and standard deviation reported.

### **α**-MSH reduces activation of cellular autophagy by Rapamycin

10,000 MNT-1 cells were seeded per well in a 96-well flat bottom tissue culture plate (Costar 3596) in 100 microliters of growth media and placed in a 37°C humidified incubator with 5% CO_2_. After 48 hours, 100nM rapamycin (AG Scientific, R1018) alone or mixed with 100nM **α**-MSH or assay buffer was added to the cells in triplicate wells in a final volume of 200 microliters. After 7 days, melanin levels were determined in each well as described above using the “Conventional method.”

### ASIP reverses autophagic impairment induced by **α**-MSH

2,000 MNT-1 cells were seeded per well in a 96-well flat bottom tissue culture plate (Costar 3596) in 100 microliters of growth media and placed in a 37°C humidified incubator with 5% CO2. After 72 hours, 100 microliters of growth media containing 40nM (final 20nM) or 400nM (final 200nM) of **α**-MSH or growth media alone was added to the cells in triplicate wells in a final volume of 200 microliters. After 2 days, media was aspirated from each well and 100 microliters of growth media containing 100nM of rhASIP or growth media alone was added to each well. After 4 days, melanin levels were determined in each well as described above using the “Alternative method.”

### Glucose level determination

200,000 MNT-1 cells were seeded in a 25 cm^2^ canted neck tissue culture flask (353109) in 5 ml of growth media containing either 200nM **α**-MSH or control buffer and placed in a 37°C humidified incubator with 5% CO_2_. The concentration of glucose in the culture medium was measured in technical duplicate using an Accu-Chek glucose meter and strips on day 0, after 5 and 6 days in culture.

### Detection of apoptosis by fluorescence microscopy

10,000 MNT-1 cells were seeded in each well of an 8-chamber glass slide (Lab-Tek, 154941) in 400 microliters growth media and placed in a 37°C humidified incubator with 5% CO_2_. After 48 hours, 200 microliters of supernatant was removed and replaced with an equal volume of growth medium containing 200nM **α**-MSH (100nM final concentration) or with growth medium alone. After 6 days, 30 microliters of the apoptosis detection reagent (CellEvent™ Caspase-3/7 Green Detection Reagent, Thermofisher, C10723) was added to each well and the slide returned to the incubator for another 60 minutes. After removal of media containing the detection reagent, the slide was mounted with ProLong® Gold antifade reagent and coverslip placed. The slide was viewed (14X and 28X magnification) using a fluorescence microscope (Olympus BH2-RFCA with BP490 cube for blue wavelengths) and the image captured using a Zeiss AxioCam MRm camera. See [Supplementary Figure 2](https://drive.google.com/file/d/1tFwqvDZiiPwewzBPok5ONSH2jMTKLrIa/view?usp=sharing) for a description of two independent experiments measuring apoptosis by flow cytometry.

### Measurement of cell viability in a 96-well plate format

2,000 MNT-1 cells were seeded per well in a 96-well flat bottom tissue culture plate (Costar 3596) in 100 microliters of growth media and placed in a 37°C humidified incubator with 5% CO_2_. After 48 hours, 50 microliters of plain DMEM containing 200nM of **α**-MSH (final 50nM) or plain DMEM alone were added to the cells in triplicate wells, followed by 50 microliters of plain DMEM containing rhASIP prepared at 800nM (4X, final 200nM), 400nM (2X, final 100nM), or 200nM (1X, final 50nM) or plain DMEM alone in a final volume of 200 microliters per well, and the plate returned to the incubator. After 6 days, media was removed from each well and replaced with 200 microliters of 1X PBS. 1X PBS was removed and the plate dabbed upside-down and allowed to sit this way over a stack of paper towels for 2 minutes. After another dab, 100 microliters of CellTiter 96® AQueous One Solution Cell Proliferation Assay solution (MTS, Promega G3580) prepared in DMEM + 5% FBS was added to each well, after which the plate was returned to the incubator with the lid off and allowed to develop. The absorbance at 490nm and 650nm (background absorbance) was measured using a plate reader (Molecular Devices Thermomax and Softmax Pro software).

### Immunofluorescence detection of alpha-synuclein accumulation in MNT-1

8,000 MNT-1 cells were seeded in each well of an 8-chamber glass slide (Lab-Tek, 154941) in 400 microliters growth media and placed in a 37°C humidified incubator with 5% CO_2_. After 72 hours, supernatant was removed and replaced with either an equal volume of growth medium containing 25nM **α**-MSH, 25nM **α**-MSH + 100nM rhASIP, or with growth medium alone. After another 48 hours in culture, supernatant was removed by aspiration and 500 microliters of fixation buffer (4% paraformaldehyde in PBS, Biolegend) added to each well and the slide kept in the dark at room temperature. After 5 hours, fixation buffer was removed and 500 microliters of 1X PBS added, and then aspirated. 1/500 diluted alpha-synuclein monoclonal antibody (Syn 211, Thermofisher, AHB0261) prepared in intracellular staining buffer (ISB, 0.25% Triton-X 100, 5% goat serum in 1X PBS) was added at 400 microliters per well and the slide place in 4°C. After two washes with ISB, 400 microliters of 1/1000 diluted anti-mouse IgG-Alexafluor 488 (Biolegend, 405319) was added and the slide returned to 4°C. After two washes with ISB and one wash with 1X PBS, the slide was mounted with ProLong® Gold antifade reagent and coverslip placed. The slides were viewed (28X magnification) using a fluorescence microscope (Olympus BH2-RFCA with BP490 cube for blue wavelengths) and the position on the slide was randomly selected and captured using a Zeiss AxioCam MRm camera (Exposure: 400 ms, 5% gain). Image analysis was done using the Image-J software (see [Supplementary Table I](https://drive.google.com/file/d/1TuqhiNMlMDRwOQZe-Wx2A4lH15EBa1FG/view?usp=sharing) for settings and images).

### Detection of MC1R in human SN

Tissue Processing: All brains were processed as described (Chu et.al., 2006). Briefly, each brain was cut into 2 cm coronal slabs and then hemisected. The slabs were fixed in 4% paraformaldehyde for 5 days at 4°C. After brain blocks were sampled from one side of the brain for pathologic diagnoses, the remaining brain slabs were cryoprotected in 0.1M phosphate buffered saline (PBS; pH 7.4) containing 2% dimethyl sulfoxide and 10% glycerol for 48 hours followed by 2% dimethyl sulfoxide and 20% glycerol in PBS for at least 2 days prior to sectioning. The fixed slabs were then cut into 18 adjacent series of 40 μm thick sections on a freezing sliding microtome. All sections were collected and stored at −20°C in a cryoprotectant solution prior to processing. Immunohistochemistry: An immunoperoxidase labeling method was used to examine the expressions of MC1R. Endogenous peroxidase was quenched by 20 minutes of incubation in 0.1M sodium periodate, and background staining was blocked by 1 hour of incubation in a solution containing 2% bovine serum albumin and 5% normal goat serum. Tissue sections were incubated in anti-MC1R rabbit polyclonal antibody (1:200; Thermofisher Cat# PA5-33923) overnight at room temperature. After 6 washes, sections were sequentially incubated for 1 hour in biotinylated goat anti-rabbit IgG (1:200; Vector), followed by Elite avidin-biotin complex (1:500; Vector) for 75 minutes. The immunohistochemical reaction was completed with 0.05% 3,3’-diaminobenzidine and 0.005% H_2_O_2_. Sections were mounted on gelatin-coated slides, dehydrated through graded alcohol, cleared in xylene, and coverslipped with Cytoseal (Richard-Allan Scientific, Kalamazoo, MI). Immunohistochemical controls: Immunohistochemical control experiments included omission of the primary antibodies (which control for the specificity of the staining procedure and the secondary antibody) and replacement of the primary antibodies with an irrelevant IgG matched for protein concentration. The control sections were processed in a manner identical to that described above. An adsorption control experiment for MC1R antibody was also done. Briefly, the anti-MC1R rabbit antibody was combined with a five-fold (by weight) of recombinant MC1R peptide (217-232, Alomone Labs) in TBS and incubated at 4°C overnight. The immune complexes with the antibody and blocking peptide were centrifuged at 10,000g for 20 min. The adsorbed peptide/antibody supernatant was then used in lieu of the primary antibody. This resulted in a total absence of staining (see Figure 5).

### Intranasal administration of **α**-MSH

The animal studies described were approved by InTouch BioSolutions LLC Institutional Animal Care and Use Committee (IACUC Protocol # 052017). Trial 1: 16-17 weeks-old C57BL/6J female mice (JAX, 000664); Trial 2: 30 weeks-old C57BL/6NTac female mice (Taconic Biosciences, Inc., B6-F) were administered 3.32 ug of **α**-MSH in 10 microliters aCSF or aCSF alone (control mice) every other day for a total of 14 intranasal administrations over 26 days (1 month). A Rainin L20 pipette fitted with a tip was used to deliver 10 microliters into the left nostril of toe-pinched confirmed fully anesthetized mouse. After administration, the mouse was allowed to remain in a supine position for 10 minutes under anesthesia and subsequently allowed to fully recover in its respective cage. Litter-mates comprised of two to three mice administered with **α**-MSH and one mouse administered with aCSF.

### Dopamine replacement and treatment with a dopamine receptor agonist

Levodopa (L-dopa, PHR1271-500MG), Benserazide hydrochloride (B7283-1G), L-ascorbic acid (A4544-25G), and Pramipexole (PHR1598-500MG) were purchased from Sigma Aldrich, St. Louis, MO., and prepared in 1X PBS (Gibco 10010-023). Dopamine replacement: Mice received intraperitoneal injection of 0.5 ml solution (2.5 mg/ml L-ascorbic acid in 1X PBS) containing 2.5 mg/ml L-dopa and 0.75 mg/ml Benserazide hydrochloride for a final L-dopa dose of approximately 50 mg/kg and Benserazide hydrochloride dose of approximately 15 mg/kg per mouse (approximate weight of 25g). Gait deficit was assessed after 3 hours of injection (“ON” state) and after 8 hours and 11 hours of injection (“OFF” state). In a separate experiment, reducing Benserazide hydrochloride by 10-fold (1.5 mg/kg) and omitting L-ascorbic acid led to similar results. Dopamine receptor agonist treatment: Mice received intraperitoneal injection of 0.2 ml solution (1X PBS) containing 0.135 mg/ml Pramipexole for a final dose of approximately 1 mg/kg per mouse (approximate weight of 27g). Gait deficit was assessed before injection (“OFF” state), after 2 hours of injection (“ON” state) and after 6 hours of injection (“OFF” state).

### Assessment of gait deficit by a modified rotarod

Gait was assessed using a Rotarod apparatus (IITC Life Science Inc. Series 8) every week until the end of the study using a modified drum consisting of a polyethylene pipe insulation of 1¼ inch diameter (Grainger, 2CKE8) to encourage walking. A digital video camera (Contour 2+) was used to record the hind legs. Mice were allowed to warm-up for 2 minutes at the top rotational speed of 30 RPM. The rotarod was quickly restarted and the run recorded for another 5 minutes at the top speed of 30 RPM (Trial 1) or 20 RPM (Trial 2, starting at week 16 after the one-month induction period).

### Computer-assisted gait analysis

We developed a computer-assisted (machine learning) gait analysis software using the DeepLabCut platform [31] to train an object tracking model. After 150K training iterations, we used the software to track the mouse paws, ankles, knees, and tailbase from recorded rotarod run videos. The mean distance (in pixel units) between objects was derived from the average of the relative maximums and minimums and the ankle joint angles (in degrees) were derived using DLC2Kinematics, a post-processing DeepLabCut module [31]. A manuscript is under preparation describing the details of the software development.

### Brain perfusion and fixation

Brains of anesthetized mice were perfused with 24ml 0.9% NaCl saline solution. Briefly, the heart was exposed and the descending aorta clamped with a bulldog clamp. Saline was administered via the left ventricle using a 22G 1-1/2 in needle, right atrium clipped with a scissor, and perfusion was terminated after 6 minutes. The brain was carefully removed and placed in 10 ml of 4% PFA in 1X PBS solution. After 3 days, the brain was washed 3Xs with 20ml of saline and placed in 1X PBS and stored in 4°C.

### Neurohistology Embedding, Sectioning & Staining and imaging/scanning

Brains were examined then treated overnight with 20% glycerol and 2% dimethylsulfoxide to prevent freeze-artifacts. The specimens were then embedded in a gelatin matrix using MultiBrain®/ MultiCord® Technology (NeuroScience Associates, Knoxville, TN). The blocks were rapidly frozen, after curing by immersion in 2-Methylbutane chilled with crushed dry ice, and mounted on a freezing stage of an AO 860 sliding microtome. The MultiBrain®/MultiCord® blocks were sectioned in coronally with 35 micrometer (µm) setting on the microtome. All sections were cut through the entire length of the specimen segment and collected sequentially into series of 24 containers. All containers had Antigen Preserve solution (50% PBS pH7.0, 50% Ethylene Glycol, 1% Polyvinyl Pyrrolidone); no sections were discarded. For immunohistochemistry: All incubation solutions from the blocking serum onward used Tris buffered saline (TBS) with Triton X-100 as the vehicle; all rinses were with TBS. After a H_2_O_2_ treatment and blocking serum, the sections were immunostained with 1/30,000 diluted rabbit anti-alpha-synuclein (phospho S129) antibody [EP1536Y] (Abcam, ab51253) and incubated overnight at room temperature. Vehicle solutions had Triton X-100 for permeabilization. Following rinses, a biotinylated secondary anti-rabbit IgG was used. After further rinses, Vector Lab’s ABC solution (avidin-biotin-HRP complex; details in instruction for VECTASTAIN® Elite ABC, Vector, Burlingame, CA) was used. The sections were again rinsed, then treated with diaminobenzidine tetrahydrochloride (DAB) with nickel and H_2_O_2_ to create a visible reaction product. Following further rinses, the sections were mounted on gelatin coated glass slides, and air dried. The slides were dehydrated in alcohols, cleared in xylene, and coverslipped. Image capture: The slides were scanned using Huron digital photography system at 20x resolution (0.4microns/pixel). For immunofluorescent histochemistry: All incubation solutions from the primary antibody onward used Tris buffered saline (TBS) with Triton X-100 as the vehicle; all rinses are with TBS. The sections were immunostained with the primary antibody: 1/500 diluted guinea pig anti-p62/SQSTM-1 (ProGen, GP62-C) and 1/1500 chicken anti-tyrosine hydroxylase (Encor, CPCA-TH) overnight at room temperature. Vehicle solutions had Triton X-100 for permeabilization. Following rinses, 1/500 diluted anti-Guinea Pig AF488 (Jackson Labs, 706-545-148) or 1/500 donkey anti-Chicken Cy3 (Jackson Labs, 703-165-155) were used. Following rinses, Hoechst (a nissl counterstain) was used. Following further rinses, the sections were mounted on gelatin coated glass slides, and air dried. The slides were dehydrated in alcohols, cleared in xylene and coverslipped. Image capture: The slides were scanned using Olympus VS200 scanning system at 20x resolution (0.274um/pixel) using the Olympus VS200 ASW software. Fluorescent channel exposure settings were as follows: DAPI (Hoechst) - 214.998, FITC (p62) - 112.999 and TRITC (TH) - 47.101.

### Fluorescence image analysis

We developed a computer-assisted histology image processing software using OpenCV [32]. The software creates a border around the population of tyrosine hydroxylase (TH+) positive dopaminergic cells within the substantia nigra pars compacta (SNpc) and ventral tegmental area (VTA) region by running the image through a series of dilations and erosions to produce a contour map. This is converted into a bitwise mask function which is used to extract the region of interest (ROI) from each fluorescence channel. A manuscript is under preparation that describes the details of the software development. Using Image-J software, mean, area and integrated density fluorescence intensity was determined for each fluorescence channel e.g., green channel for p62, within the ROI.

### Statistical analysis

Information can be found in the figure or table legends.

## Results

We chose a melanized human melanoma cell line, MNT-1 [33], to model SN DNs for the following reasons. Melanoma cells and SN DNs share a common embryonic origin (neural crest), are pigmented and express alpha-synuclein [34]. Moreover, a clear association between melanoma and PD incidence [35, 36] suggested the possibility of a shared etiology involving the loss of autophagy in malignancy [37, 38] and in dopaminergic dysfunction, as mentioned. Furthermore, both melanoma cells and SN DNs produce dopamine [39].

Neuromelanin is a product of autophagic reactions in the SN [40]. Indeed, decreased neuromelanin in PD coincided with “autophagic degeneration” in melanized neurons of the SN in PD patients [6, 40]. Melanin levels correlate with activation of autophagy in MNT-1 [41, 42]. Melanin formation induced by autophagy activators was significantly reduced in the presence of **α**-MSH (Figure 1, [Supplementary Figure 1](https://drive.google.com/file/d/1s5mTF8NlumnD4xRiTB2gxRy6hcKeUfRP/view?usp=sharing)). In addition, a brief exposure to **α**-MSH was sufficient to impair cellular autophagy in MNT-1, such that replacing the culture medium without **α**-MSH did not restore autophagy-dependent melanin formation (Figure 2). On the other hand, replacing the medium with one containing ASIP, a natural biologic inhibitor of **α**-MSH [43, 44], reversed autophagic impairment induced by **α**-MSH and melanin formation was restored (Figure 2). As an inverse agonist, ASIP can engage the MC1R receptor and induce the opposite signal (e.g., reduction of intracellular cAMP levels), thereby effectively reversing the effects of **α**-MSH [44–47].

**Figure 1:**
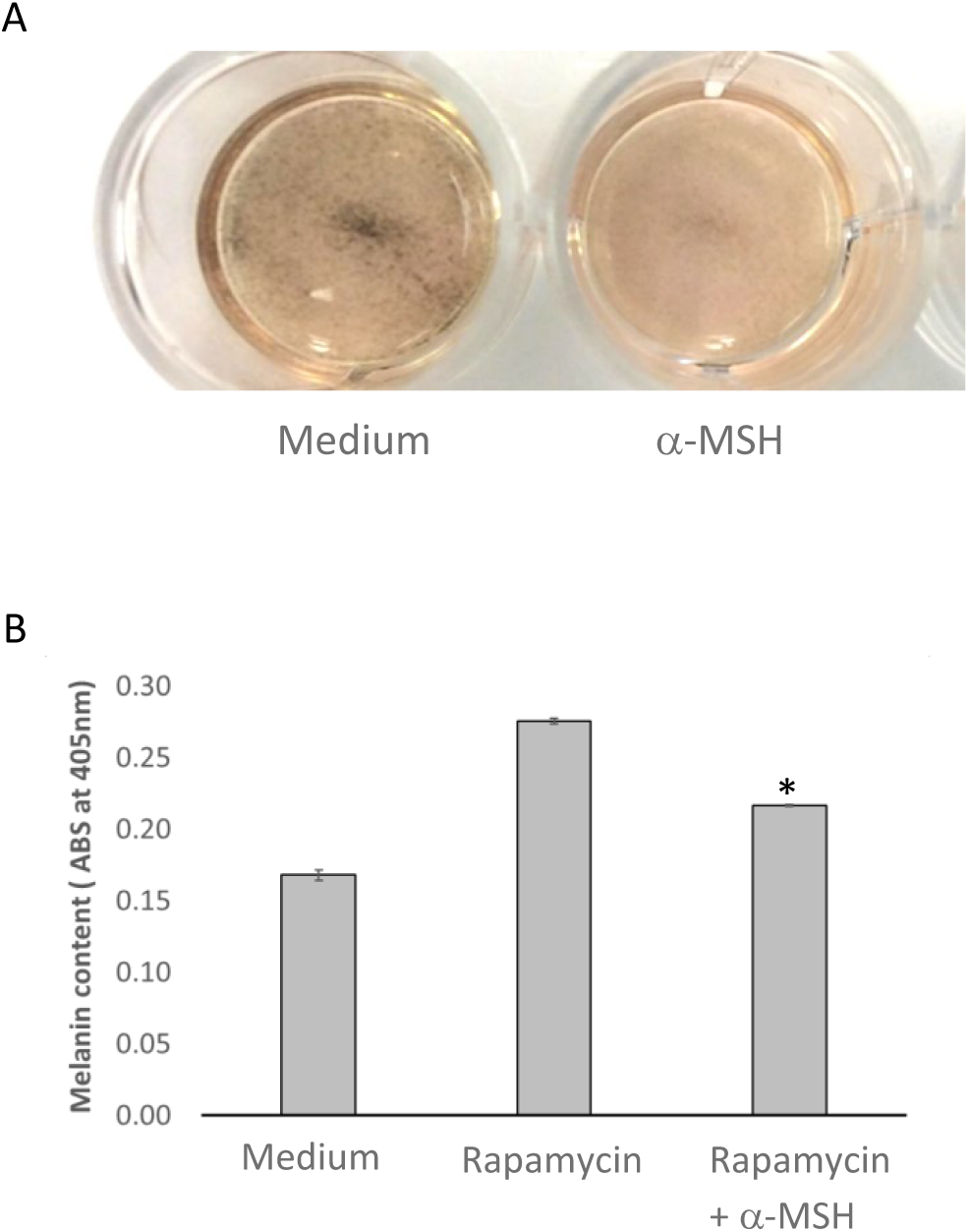
*α*−MSH reduces autophagy-dependent melanin formation in MNT-1 cells. (A) Starvation stimulates autophagy-dependent melanin formation in MNT-1 cells. MNT-1 cells were allowed to remain in culture for 9 days to starve and stimulate autophagy-dependent melanin formation (left well, dark substance). Melanin formation was reduced in starved MNT-1 cells in the presence of *α*−MSH (right well, diminished dark substance), suggesting inhibition of starvation-induced cellular autophagy. (B) The autophagy activator, rapamycin, stimulates autophagy-dependent melanin formation in MNT-1 cells. Reduced melanin formation is observed in the presence of *α*−MSH, suggesting inhibition of rapamycin-induced cellular autophagy. Error bars are SD of mean of triplicate measurements. *Rapamycin vs. Rapamycin + *α*−MSH, unpaired Student’s T-test (*p* value = 0.0001). A *p* value of less than 0.05 is considered significantly different. See Materials and Methods for experiment details.

**Figure 2:**
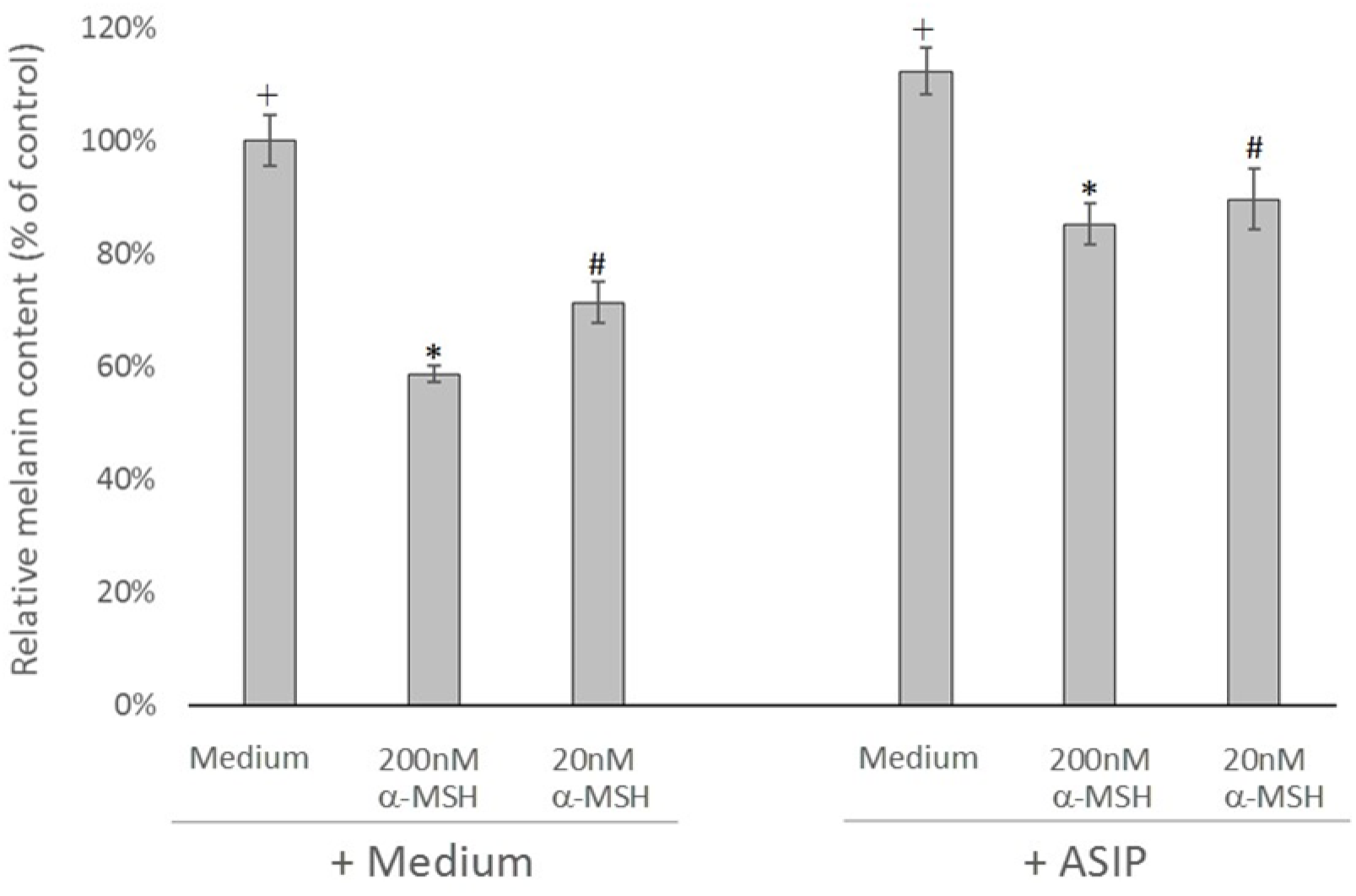
*α*−MSH induced impairment of cellular autophagy is reversed after treatment with ASIP. ASIP (acting as inverse agonist) reverses *α*−MSH effects and restores autophagy-dependent melanin formation. Cells are cultured in either medium alone or medium containing 200nM or 20nM *α*−MSH. After 2 days, media is replaced with medium alone or ASIP (100nM) and melanin levels are measured after 4 days using a fluorescence-quenching method (see Materials and Methods for experiment details). Error bars are SD of mean of triplicate measurements. “Control” is MNT-1 cells cultured in medium alone and the media replaced with medium alone. Unpaired Student’s T-test analyses of means and *p* values: ^+^ Medium vs. Medium/ASIP, *p* = 0.03; * 200nM *α*−MSH vs. 200nM *α*−MSH/ASIP, *p* = 0.0002; ^#^ 20nM *α*−MSH vs. 20nM *α*−MSH/ASIP, *p* value = 0.007. A *p* value of less than 0.05 is considered significantly different.

When nutrients are low, the mTOR signaling pathway is inhibited and cellular autophagy is activated, resulting in a decrease in the rate of consumption and slower glucose metabolism [48, 49]. Conversely, cellular autophagy is less active or inhibited when nutrient levels are adequate, allowing consumption of nutrients to proceed at a higher rate [49]. **α**-MSH activates the mTOR signaling pathway following MC1R engagement [50] and stimulates glycolysis and glucose uptake [51]. Similarly, MNT-1 cells cultured in the presence **α**-MSH did not respond to diminishing nutrients in the media and consumed glucose at a much higher rate than untreated cells (Figure 3a). Over time, the cells became apoptotic, presumably due to glucose deprivation(Figure 3b, [Supplementary Figure 2](https://drive.google.com/file/d/1tFwqvDZiiPwewzBPok5ONSH2jMTKLrIa/view?usp=sharing)). ASIP neutralized the effects of **α**-MSH in a dose dependent fashion, which allowed the cells to enter autophagy and remain viable (Figure 3c).

**Figure 3:**
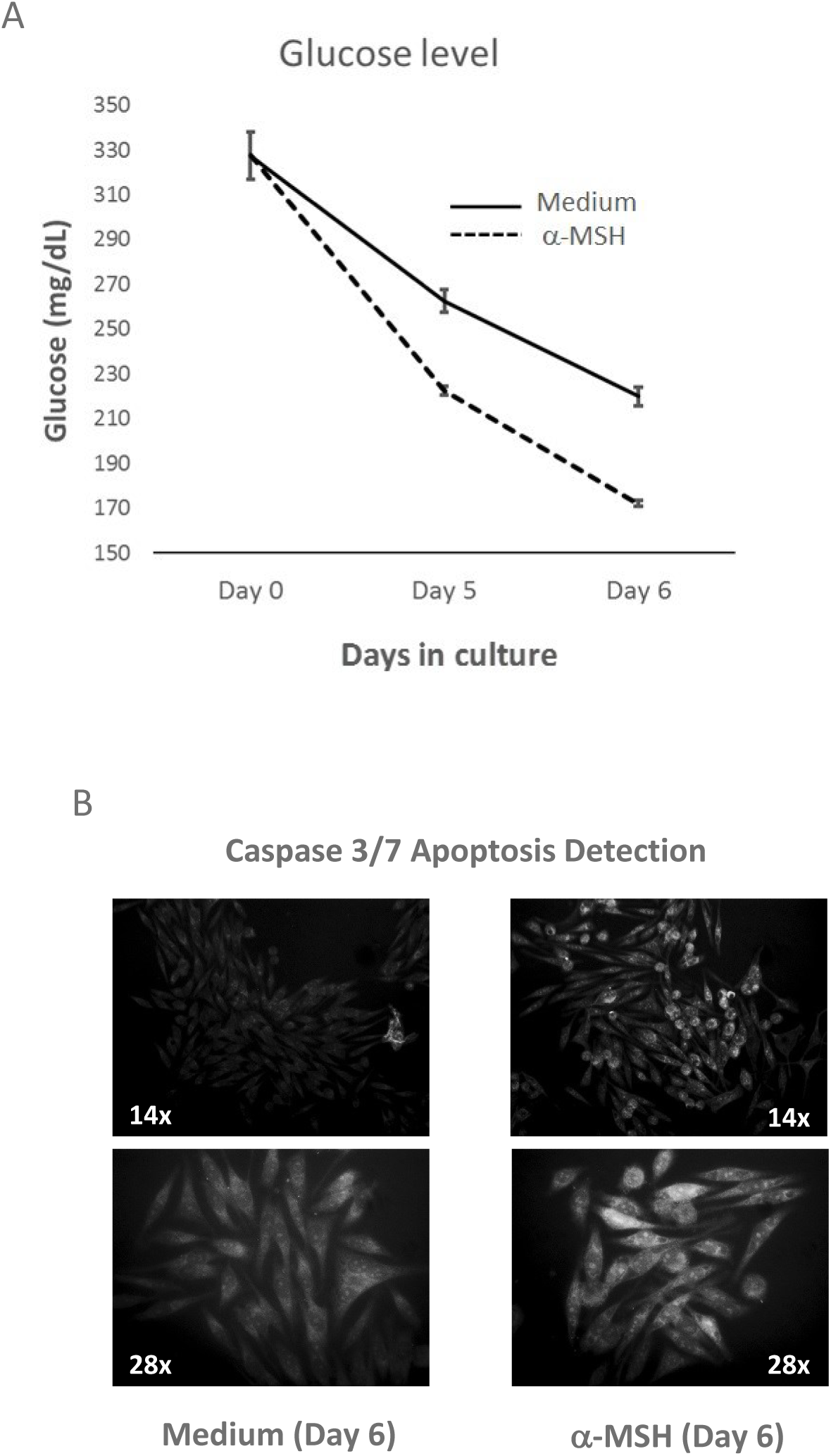
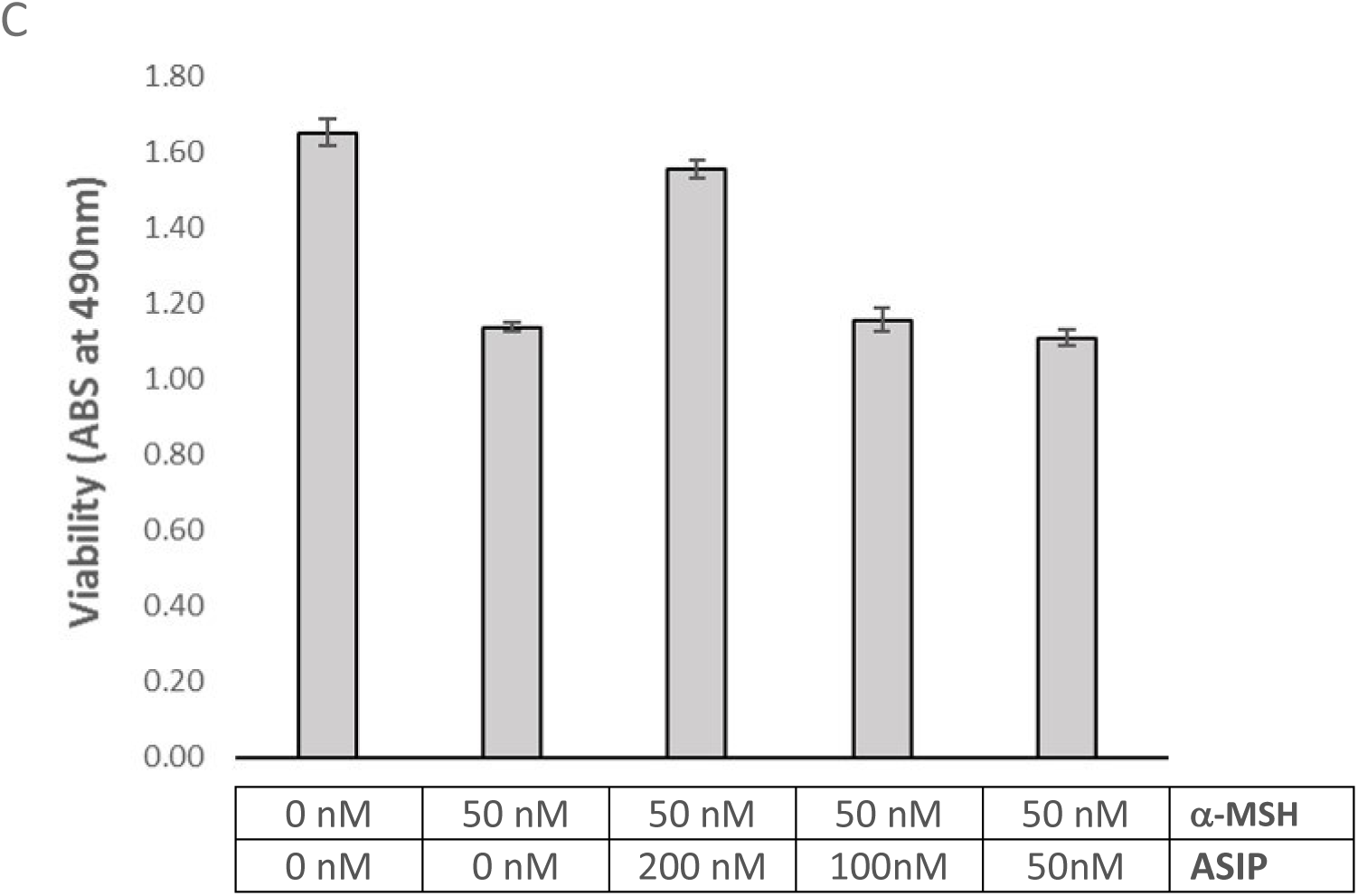
MNT-1 cells become hypermetabolic in the presence of *α*−MSH, quickly consuming glucose in culture and subsequently die by apoptosis; ASIP blocks the effects of *α*−MSH and prevents cell death. (A) *α*−MSH-treated cells consumed glucose faster than untreated cells (Medium). Error bars are SD of mean of technical duplicate measurements. (B) After 6-days, *α*−MSH-treated cells (right photos) appeared apoptotic after incubation with the Caspase 3/7 apoptosis detection reagent, which generates a bright green fluorescence in apoptotic cells with activated caspase-3/7. Untreated cells (left photos) remained non-apoptotic and healthy. Similar results were observed by flow cytometry ([Supplementary Figure 2](https://drive.google.com/file/d/1tFwqvDZiiPwewzBPok5ONSH2jMTKLrIa/view?usp=sharing)). (C) ASIP blocked the effects of *α*−MSH and protected against cell death in a dose-dependent fashion after 6-days in culture. Error bars are SD of mean of triplicate measurements.

Accumulation of alpha-synuclein is observed in PD post-mortem nigral tissue and impaired autophagy in PD has been linked to higher levels of alpha-synuclein [52, 53]. Therefore, since **α**-MSH impaired autophagy in MNT-1 (Figures 1 and 2), we tested whether accumulation of alpha-synuclein may be observed in MNT-1 cells treated with **α**-MSH. We observed robust accumulation of alpha-synuclein in MNT-1 cells cultured in the presence of **α**-MSH as compared to untreated cells. As expected, ASIP blocked the effects of **α**-MSH and inhibited alpha-synuclein accumulation (Figure 4, [Supplementary Table I](https://drive.google.com/file/d/1TuqhiNMlMDRwOQZe-Wx2A4lH15EBa1FG/view?usp=sharing)).

**Figure 4:**
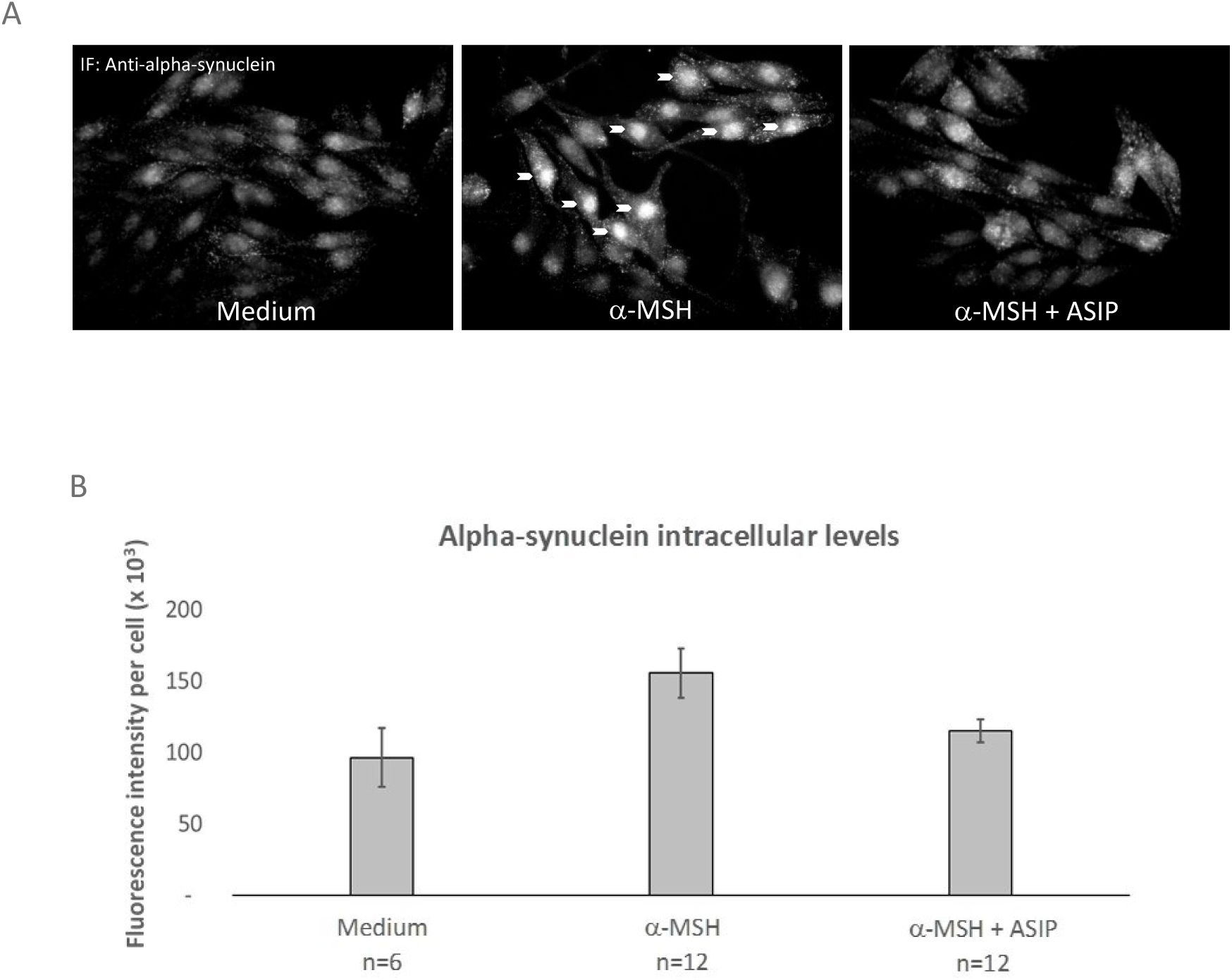
Accumulation of alpha-synuclein in MNT-1 cells is induced by *α*−MSH. (A) Representative immunofluorescent images (28X magnification) showing MNT-1 cells in medium alone, with *α*−MSH, or with *α*−MSH plus ASIP, stained with anti-synuclein antibody (Syn 211). Increased fluorescence staining, or accumulation of alpha-synuclein (arrows) was observed in the cells incubated with *α*−MSH (middle photo) as compared to cells incubated with medium alone (left photo). ASIP blocked *α*−MSH induced accumulation of alpha-synuclein (right photo). (B) A total of 30 images were processed using the Image-J analysis software and the fluorescence integrated density measured and divided by the number of cells in the image. The mean was calculated from several (n) images and reported. Error bars are SEM of mean of replicate measurements. Student’s T-test analysis of means and *p* value (two-tailed): *α*−MSH (n=12) vs. *α*−MSH + ASIP (n=12), *p* < 0.05; Welch’s T-test analysis of means and *p* values (two-tailed): Medium (n=6) vs. *α*−MSH (n=12), *p* < 0.05 and Medium (n=6) vs. *α*−MSH + ASIP (n=12), *p* = 0.4. A *p* value of less than 0.05 is considered significantly different. See Materials and Methods for experiment details and [Supplementary Table I](https://drive.google.com/file/d/1TuqhiNMlMDRwOQZe-Wx2A4lH15EBa1FG/view?usp=sharing), to view additional images and values used to calculate the means.

The MC1R receptor for **α**-MSH is expressed in the human (Figure 5 and [54]), in the non-human primate (data not shown) and in the mouse SN DNs [55]. To replicate the conditions observed in PD [29, 30] and determine if a causal relationship between elevated **α**-MSH in the brain and dopaminergic dysfunction could be observed *in-vivo*, we delivered **α**-MSH (1.6 kDa) to the brain of mice by intranasal route (Figure 6a). This method is non-invasive [56–59] and is capable of delivering much larger molecules (e.g., up to 150 kDa antibodies in rodents) to the brain [59, 60]. More importantly, this route targets the basal ganglia components, which include the SN [59, 60].

**Figure 5:**
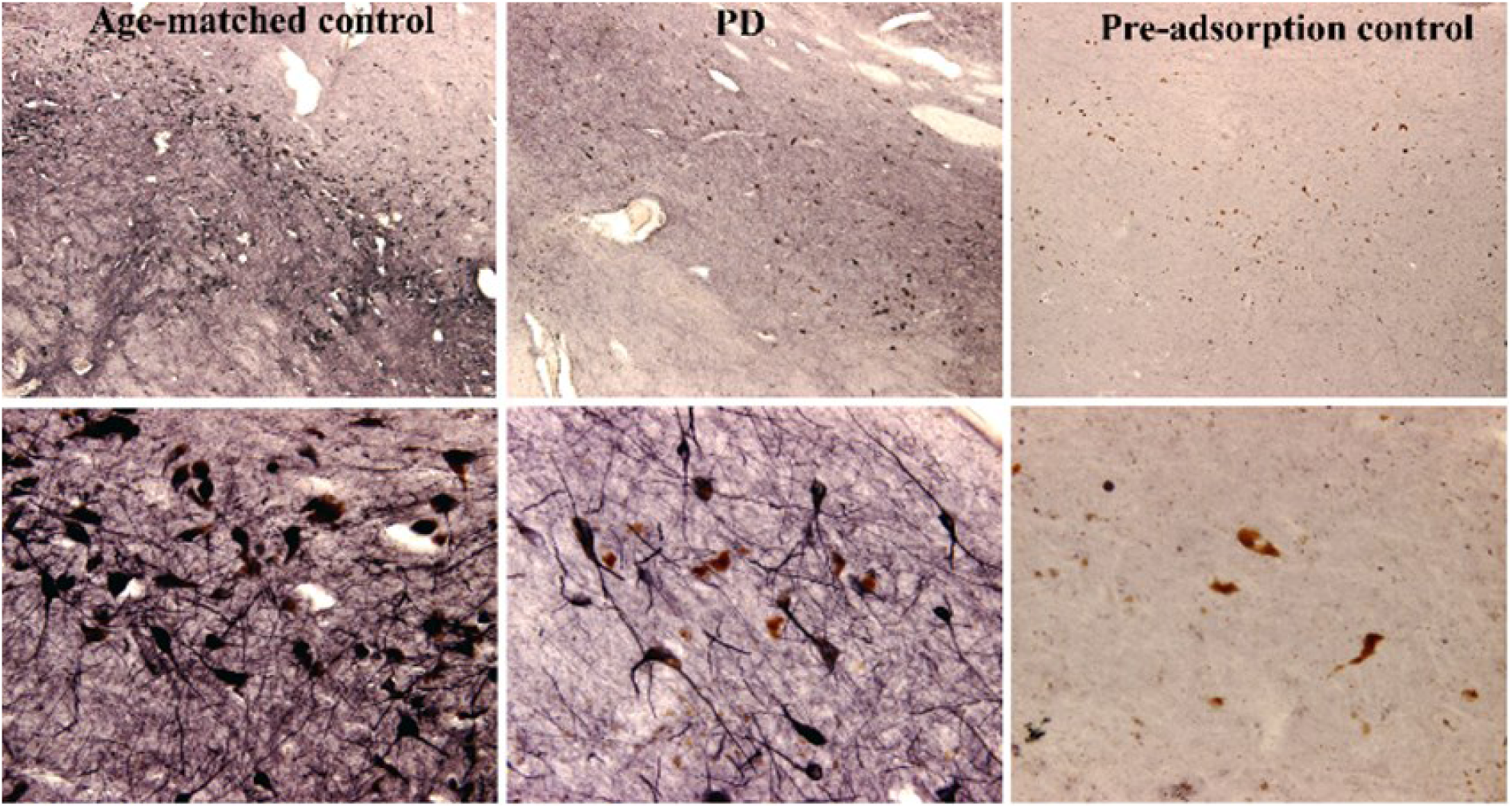
MC1R is expressed in dopaminergic neurons in the substantia nigra. Top (low magnification) and bottom (high magnification) immunohistochemistry images of anti-MC1R staining of human SN neurons from age-matched control (left) and Parkinson’s disease (PD, center) sections. MC1R staining was confirmed by MC1R peptide fragment pre-adsorbed anti-MC1R antibody (right). See Materials and Methods for experiment details.

**Figure 6:**
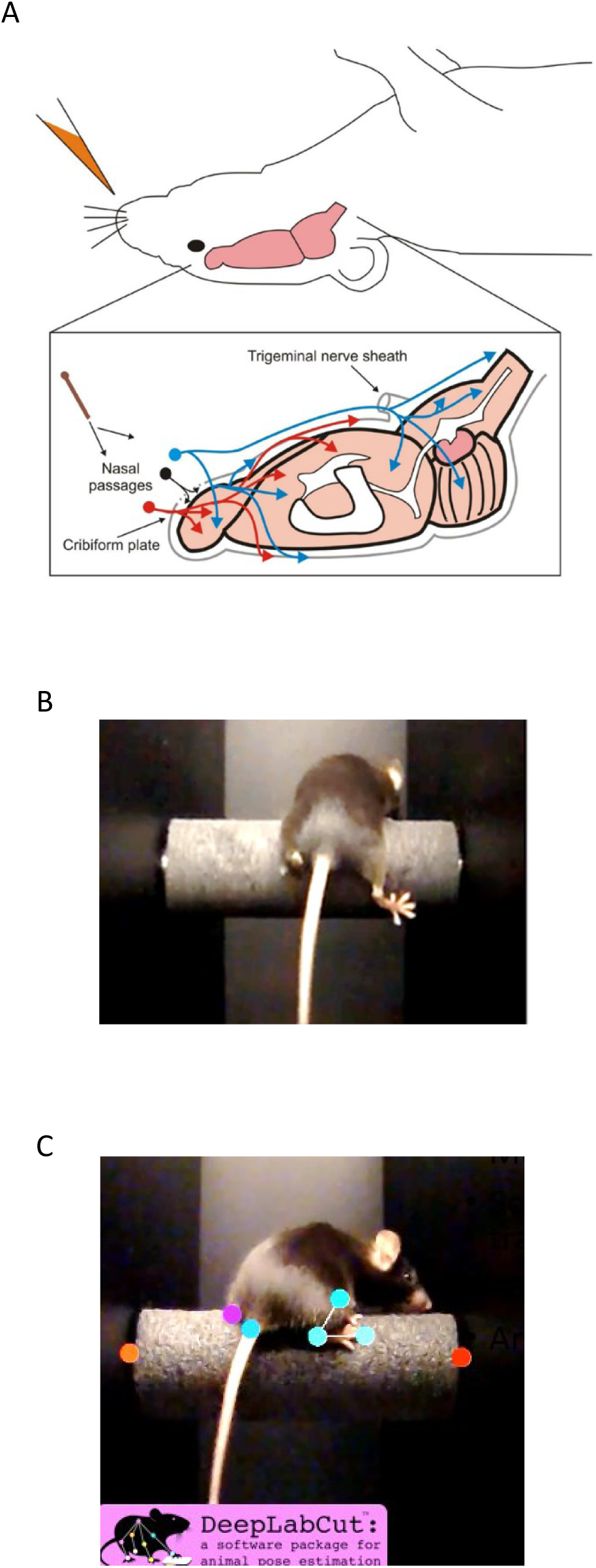
Intranasal administration, gait assessment on a modified rotarod and gait analysis by an object tracking software. (A) This figure from Malhotra et al., 2014 is used to illustrate intranasal administration of aCSF or *α*−MSH into the nostril of a mouse and uptake and route of delivery into the brain via olfactory nerves (in red) and trigeminal nerves (in blue). (B) A modified rotarod is shown on which mouse gait was recorded using a video camera. (C) A machine learning software was used for markerless tracking of objects and analysis of mouse gait and movement on the rod (see [Supplementary Video 1](https://drive.google.com/file/d/1SP0ngmMquihkBPMfnOwefMPA_CAFYOxL/view?usp=sharing)).

To maintain CNS exposure to elevated **α**-MSH, mice were administered **α**-MSH in the left nostril once every other day for a period of one month (Fig 6a). After the induction period, motor deficits were measured during the subsequent two months (Trial 1, pilot study) or six months (Trial 2). The two trials were done over two years apart in different settings, and the mice were handled by different technicians using the same strain of female mice obtained from different vendors. These differences helped to ensure that animal behavior and outcomes would be independent of variables associated with their surroundings and handling.

We used a modified rotarod designed specifically to assess gait abnormalities (Figure 6b). Rotarod runs were video recorded weekly and gait deficits inspected visually and with the use of a markerless object tracking software [31], which was configured to perform unbiased accurate measurements of the distance between the paws and the base of the tail and the ankle joint angles (Figure 6c, [Supplementary Video 1](https://drive.google.com/file/d/1SP0ngmMquihkBPMfnOwefMPA_CAFYOxL/view?usp=sharing)). To monitor the progression of motor deficit for each mouse, we established a scoring system based on movement signatures derived from these parameters and scored for rigidity as seen in the shift in the center-of-gravity (asymmetric gait), leg dragging, and loss of ankle joint angle flexion and freedom of movement.

In the first trial, three C57BL/6 female mice received intranasal **α**-MSH dissolved in artificial cerebrospinal fluid (aCSF) and one mouse received aCSF alone and served as a control. One to two weeks after the one-month induction period, a right-ward shift in the center-of-gravity began to appear in the gait of mice administered with **α**-MSH, while the control mouse remained with normal gait (Table I). At the end of the trial (Week 8), mice administered **α**-MSH had asymmetric gait as seen in the increased percentage difference between left and right ankle joint angles (Table I). The right-shifted center-of-gravity in the three mice indicated a right-side impairment and was seen in the shift of the weight-bearing left leg inward, accompanied by a corresponding decrease in the angle of the leg’s ankle joint (Table I), while the affected right leg extended outward and appeared rigid during the run (see Figure 6b, [Supplementary Video 2](https://drive.google.com/file/d/1d9BS1Aw1jNj_Ynnrjox2Qf1KK0TltVsQ/view?usp=sharing), [Supplementary Video 3](https://drive.google.com/file/d/161dcaztb__K7k5SqYJh8Eo4WQvd_FQUc/view?usp=sharing) and [Supplementary Video 4](https://drive.google.com/file/d/1Ghgz-Rxz4VTxrWextkhlmxqdyJ9B2yGx/view?usp=sharing)). The gait of the control mouse remained normal and symmetric (Table I, [Supplementary Video 5](https://drive.google.com/file/d/1n7P1i6cCJgNymI5EXxtExPRullxHvap1/view?usp=sharing)).

**Table I.** Progressive loss of gait symmetry in mice administered *α*−MSH (Trial 1) After one month of intranasal administration of *α*−MSH (n=3 mice) or artificial cerebrospinal fluid (aCSF, n=1 control mouse), rotarod run was video recorded weekly until the end of the study (week 8). By week 8, except for the control mouse, asymmetric gait was observed in mice administered *α*−MSH. The left ankle angle at week 8 was smaller than measurements at week 1, while the right ankle angle remained relatively unchanged or larger (for A4) between week 1 and week 8. Moreover, asymmetric gait can be seen in the higher % difference between the left and right ankle joint angles in mice administered *α*−MSH as compared to the control mouse, which had symmetric gait. See Materials and Methods for experiment details and [Supplementary Video 2](https://drive.google.com/file/d/1d9BS1Aw1jNj_Ynnrjox2Qf1KK0TltVsQ/view?usp=sharing), [Supplementary Video 3](https://drive.google.com/file/d/161dcaztb__K7k5SqYJh8Eo4WQvd_FQUc/view?usp=sharing), [Supplementary Video 4](https://drive.google.com/file/d/1Ghgz-Rxz4VTxrWextkhlmxqdyJ9B2yGx/view?usp=sharing) and [Supplementary Video 5](https://drive.google.com/file/d/1n7P1i6cCJgNymI5EXxtExPRullxHvap1/view?usp=sharing) to view mouse A1, A3, A4 and control mouse A2, respectively.

| Mouse ID | Week post induction | Left ankle angle (°) | % change (Wk 1 vs. 8) | Right ankle angle (°) | % change (Wk 1 vs. 8) | % difference (L vs. R) ankle angles (°) at week 8 |
| --- | --- | --- | --- | --- | --- | --- |
| A2 (aCSF) | Week 1 | 103 | 0% | 104 | 3% | 5% |
|  | Week 8 | 103 |  | 108 |  |  |
| A1 (α-MSH) | Week 1 | 99 | 7% | 100 | 1% | 9% |
|  | Week 8 | 93 |  | 101 |  |  |
| A3 (α-MSH) | Week 1 | 98 | 11% | 105 | 3% | 23% |
|  | Week 8 | 87 |  | 109 |  |  |
| A4 (α-MSH) | Week 1 | 106 | 15% | 90 | 26% | 23% |
|  | Week 8 | 90 |  | 113 |  |  |

Having observed 100% of the mice developed abnormal gait after induction with intranasal **α**-MSH, we ran a second larger and longer trial and used the same dosing route and schedule as previous. The second trial included two sets of experiments. Trial 2a confirmed the reproducibility of the induction protocol and expanded the histopathology assessment to include the evaluation of autophagic dysfunction in dopaminergic neurons. Trial 2b confirmed the presence of dopaminergic dysfunction and dopamine deficiency in affected mice.

In Trial 2a, the number of mice was increased to n=6 in the **α**-MSH group and to n=3 in the aCSF group. The rotarod runs were video recorded weekly for 24 weeks (6 months). Similar to the previous observations, 2 out of 6 **α**-MSH administered mice developed gait abnormalities within 2 weeks after the one month of induction with **α**-MSH and showed a right-shifted center-of-gravity. However, these two “early onset” mice were not able to keep up with the rotational speed. Seeing that gait evaluations could proceed at a lower speed, the remaining rotarod runs, beginning on week 16 to the end of the study (week 24), were switched from 30 rpm to 20 rpm for all the mice.

All mice that received **α**-MSH developed gait abnormalities at different times, which gradually worsened (Table II). The control mice had normal gait ([Supplementary Video 6](https://drive.google.com/file/d/1gKyPdByOPbZBVwBpE7zttzteAmNzYBgS/view?usp=sharing)), with the exception of mouse A3 that developed a subtle right leg gait abnormality near the end of the trial and its scores were used as basal control values. Whereas in the previous trial, all three mice administered **α**-MSH developed a right-sided impairment, mice in this trial developed gait deficits on both sides (Table II). The most common abnormalities observed were asymmetric gait and leg drag. Leg drag was sometimes accompanied by a decrease in ankle joint flexion, which was accompanied by a decrease in the freedom-of-movement (FOM) at that ankle joint (Table IIb). These deficits reflect rigidity in the affected limb(s), including the inability of the leg to pivot medially ([Supplementary Video 7](https://drive.google.com/file/d/1bZqG_mj3-5kAIFiYKrVE-NewvMlVMWLw/view?usp=sharing), [Supplementary Video 8](https://drive.google.com/file/d/1e1tam5SbYJNDNMa6vWGgxFvQxliPcZ79/view?usp=sharing) and [Supplementary Video 9](https://drive.google.com/file/d/1Ihodmi4X-_RKz8vzqtb3JGpSI6yn3y61/view?usp=sharing)).

**Table II.** Progressive loss of gait symmetry and manifestations of rigidity in mice administered *α*−MSH (Trial 2a) (A) After one month of intranasal administration of *α*−MSH (n=6 mice, bold text) or artificial cerebrospinal fluid (aCSF, n=3 control mice), rotarod runs were video recorded weekly until the end of the study (week 24). Progressive worsening of motor deficit between week 16 and week 24 was observed in mice administered *α*−MSH that developed either a right-shifted or left-shifted center-of-gravity (CoG, loss of gait symmetry), leg drag in one or both sides, and decreased flexion and freedom-of-movement (FOM) at the ankle joints. Control mice showed no progressive motor deficit except for mouse A3, which had a slight right leg gait deficit from an unknown or natural cause. (B) A summary of features of rigidity observed for each mouse. See Materials and Methods for experiment details and [Supplementary Video 6](https://drive.google.com/file/d/1gKyPdByOPbZBVwBpE7zttzteAmNzYBgS/view?usp=sharing), [Supplementary Video 7](https://drive.google.com/file/d/1bZqG_mj3-5kAIFiYKrVE-NewvMlVMWLw/view?usp=sharing), [Supplementary Video 8](https://drive.google.com/file/d/1e1tam5SbYJNDNMa6vWGgxFvQxliPcZ79/view?usp=sharing) and [Supplementary Video 9](https://drive.google.com/file/d/1Ihodmi4X-_RKz8vzqtb3JGpSI6yn3y61/view?usp=sharing) to watch C1, A2, B1 and C3, respectively. **Footnotes:** ^1^ [Induction, side] 30 weeks-old female C57Bl/6 mice received intranasal (INS) administration of *α*−MSH (n=6) or aCSF (n=3, normal control) in the left nostril every other day for 4 weeks. ^2^ [Gait deficit onset] At the end of the one month induction period, walking was recorded on a modified rotarod and motor deficits inspected visually and by a machine learning object detection and analysis software. The time of appearance of abnormal gait was noted and reported as onset of gait deficit. ^3^ [CoG score] A shift in the center-of-gravity (CoG) indicates loss of gait symmetry. The score is a comparison of measurements from two time points (e.g., week 16 vs. week 24), from which the % change in the distance of the paws from midline and the ankle joint angles are determined and added to produce a score. (+) indicates a right-shifted CoG and (-) indicates a left-shifted CoG. ^4^ [Leg drag score] The leg drag score is a comparison of measurements from two time points (e.g., week 16 vs. week 24), from which the % change in ankle Min (minimum value) and Max (maximum value) joint angles are determined and added to produce a score. ^5^ [Decrease in ankle angle flexion] This is a comparison of measurements from two time points (e.g., week 16 vs. week 24) and the % change in the ankle joint angle Min measurements is determined for each leg. ^6^ [Decrease in ankle angle freedom-of-movement (FOM)] This is a comparison of measurements from two time points (e.g., week 16 vs. week 24) and the % change in the difference between ankle joint angles (Min and Max) is determined for each leg. ^a^ This mouse developed a right-side gait deficit prior to week 16, however, measurements at the lower rotarod speed (20 rpm) was not taken at earlier points and could not be compared with measurements taken at week 16. Therefore, the apparent lower CoG score and lower right side impairment vs. left side impairment reflects a right side that was already impaired by week 16, showing only slight worsening by week 24. ^b^ This mouse had developed a left-side gait deficit prior to week 16, as seen by the dragging of the left leg with the paw digits fanning out during down stride movement; and the inability of the paw to pivot medially and assume normal gait. However, measurements at the lower rotarod speed (20 rpm) was not taken at earlier points and could not be compared with measurements taken at week 16 when the deficit was still apparent ([Supplementary Video 9](https://drive.google.com/file/d/1Ihodmi4X-_RKz8vzqtb3JGpSI6yn3y61/view?usp=sharing)). Additionally, this mouse developed a right-side impairment starting at week 16, which progressively worsened by week 24 and quantified as shown ([Supplementary Video 9](https://drive.google.com/file/d/1Ihodmi4X-_RKz8vzqtb3JGpSI6yn3y61/view?usp=sharing)). We surmised that the impaired left leg adjusted to accommodate a right-side impairment and as a result, left leg progressive worsening was not detected between weeks 16 and 24. By visual inspection, at week 24 the mouse showed unsteady gait, with both legs appearing impaired. See [Supplementary Video 9](https://drive.google.com/file/d/1Ihodmi4X-_RKz8vzqtb3JGpSI6yn3y61/view?usp=sharing) to watch mouse C3.

| Progressive gait impairment measured between weeks 16 and 24 |  |  |  |  |  |  |
| --- | --- | --- | --- | --- | --- | --- |
| Mouse ID | Induction, side<sup>1</sup> | Gait deficit onset<sup>2</sup> | CoG score<sup>3</sup> | Leg drag score<sup>4</sup> | Decrease in ankle angle flexion<sup>5</sup> | Decrease in ankle angle FOM<sup>6</sup> |
| A3 | aCSF, left | ND | + 2 | (L) 0 , (R) 4 | (L) ND , (R) 3% | (L) ND , (R) 4% |
| A1 | α-MSH, left | Week 2 | + 9 | (L) 23 , (R) 0<sup>a</sup> | (L) 31% , (R) 6%<sup>a</sup> | (L) 32% , (R) 24%<sup>a</sup> |
| A2 | α-MSH, left | Week 1 | + 22 | (L) 0 , (R) 0 | (L) 0% , (R) 0% | (L) 0% , (R) 0% |
| B2 | aCSF, left | ND | 0 | (L) 0 , (R) 0 | (L) 0% , (R) 0% | (L) 1% , (R) 12% |
| B1 | a-MSH, left | Week 20 | + 46 | (L) 0 , (R) 0 | (L) 0% , (R) 0% | (L) 0% , (R) 0% |
| B3 | a-MSH, left | Week 12 | - 47 | (L) 10 , (R) 6 | (L) 0% , (R) 0% | (L) 0% , (R) 0% |
| C1 | aCSF, left | ND | 0 | (L) 0 , (R) 0 | (L) 0% , (R) 0% | (L) 0% , (R) 0% |
| C2 | a-MSH, left | Week 6 | 0 | (L) 15 , (R) 0 | (L) 7% , (R) 0% | (L) 0% , (R) 0% |
| C3 | a-MSH, left | Week 12 | 0 | (L) 0<sup>b</sup> , (R) 23 | (L) 0%<sup>b</sup> , (R) 27% | (L) 0%<sup>b</sup> , (R) 32% |

| Mouse ID | Impaired side(s) | Description of Impairment |
| --- | --- | --- |
| A3 | Right<sup>#</sup> | <sup>#</sup> A slight gait abnormality detected in the right leg due to unknown or natural cause. |
| A1 | Left and Right | Early onset rigidity in both legs. Progressive decline observed in right-ward shift in the center-of-gravity (CoG) with drag in the left leg. Decreased flexion and decreased freedom-of-movement (FOM) in both legs. |
| A2 | Right | Early onset rigidity in the right leg. Progressive decline observed in right-ward shift in CoG. |
| B2 | Neither | None. Reduction in right leg FOM cannot be confirmed with other deficits. |
| B1 | Right | Rigidity in the right leg. Progressive decline observed in right-ward shift in CoG. |
| B3 | Left and Right | Rigidity in both legs. Progressive decline observed in right-ward shift in CoG with drag in both legs. |
| C1 | Neither | None. |
| C2 | Left | Rigidity in the left leg. Progressive decline observed with drag and decreased flexion in the left leg. |
| C3 | Left and Right | Rigidity in both<sup>b</sup> legs. Progressive decline observed in right-ward shift in CoG with drag, decreased flexion and decreased FOM in the right leg. |

In Trial 2b, we tested the possibility of dopamine deficiency as the cause of movement deficits observed in mice administered with intranasal **α**-MSH. Similar to a drug challenge test that increases dopamine activity in the brain to alleviate motor symptoms and aid in confirming a PD diagnosis [61], three affected mice were treated with mainstay drugs for PD, using L-dopa/Benserazide or Pramipexole, a dopamine receptor agonist. Applying the same scoring system described above, gait deficits were scored based on the extent of reduction of the deficit when the drug was on board (Table III). Gait deficits were temporarily improved or reversed with dopamine replacement or agonist treatment, confirming dopaminergic dysfunction and dopamine deficiency as the driver of gait deficits observed in these mice (Table III).

**Table III.** Detection of dopaminergic dysfunction in *α*−MSH administered mice using dopamine replacement and receptor agonist treatment (Trial 2b) (A) Four to five months after the one-month induction with intranasal *α*−MSH, mice (n=3) showed asymmetric gait, leg drag in one or both sides, and with decreased flexion and decreased freedom-of-movement (FOM) at the ankle joints. Alleviation of gait deficit was observed after dopamine replacement with L-dopa/Benserazide or treatment with a dopamine receptor agonist, Pramipexole, and the degree of alleviation was scored using measurements obtained after 2-3 hours of treatment (see Table II footnotes for additional details), when the drug was on board, and measurements obtained before treatment or 6-11 hours after treatment, when the drug was no longer on board. (B) A summary of features of gait impairment observed in each mouse as revealed after dopamine replacement or treatment with agonist. See Materials and Methods for experiment details and [Supplementary Video 10](https://drive.google.com/file/d/1dc7kGG-1JTrNx3HOOUC4gghOmHs_NDwX/view?usp=sharing) (L-dopa/Benserazide) and [Supplementary Video 11](https://drive.google.com/file/d/1mzfehtT0xv-TSScSPOSVQ59NzV5in7H1/view?usp=sharing) (Pramipexole) to watch mouse 1.

| ID | Induction, intranasal side | Dopamine replacement and receptor agonist (Rx) | Shift in center of gravity (Score of <2, no shift*) | Leg drag (Score of <4, no drag*) | Decrease in ankle angle flexion (<3%, not present*) | Decrease in ankle angle FOM (<4%, no loss*) |
| --- | --- | --- | --- | --- | --- | --- |
| 1 | α-MSH, left | L-dopa/Benserazide | 39 right-shifted | (L) 0 , (R) 0 | (L) 5% , (R) 1% | (L) 12% , (R) 16% |
|  |  | Pramipexole | 24 right-shifted | (L) 6 , (R) 9 | (L) 11% , (R) 12% | (L) 18% , (R) 14% |
| 2 | α-MSH, left | L-dopa/Benserazide | 11 right-shifted | (L) 0 , (R) 13 | (L) 3% , (R) 11% | (L) 13% , (R) 8% |
|  |  | Pramipexole | 20 right-shifted | (L) 5 , (R) 42 | (L) 3% , (R) 31% | (L) 0% , (R) 0% |
| 3 | α-MSH, left | L-dopa/Benserazide | 34 left-shifted | (L) 0 , (R) 9 | (L) 0% , (R) 8% | (L) 0% , (R) 0% |
|  |  | Pramipexole | 34 left-shifted | (L) 22 , (R) 18 | (L) 17% , (R) 13% | (L) 0% , (R) 8% |

\* Base values obtained from mouse A3 (Trial 2a). See Table IIa.

| ID | Impaired side | Features of gait deficits detected |
| --- | --- | --- |
| 1 | Right dominant | Right-shifted center-of-gravity with a slight right leg drag<br>Poor left and right ankle joint flexion and freedom-of-movement (FOM) |
| 2 | Right dominant | Right-shifted center-of-gravity with right leg drag<br>Poor right ankle joint flexion and reduced left and right ankle joint FOM |
| 3 | Left dominant | Left-shifted center-of-gravity with left and right leg drag<br>Poor left and right ankle joint flexion and reduced right ankle joint FOM |

Post-mortem analysis of brains from PD patients invariably show end-stage pathologies including the loss of DNs in the SN and ventral tegmental area (VTA), reduction in striatal TH+ density and the presence of phosphorylated alpha-synuclein (pSer129+) aggregates in the form of Lewy bodies or inclusions [62, 63]. Therefore, we also looked for these pathologies in mice that developed motor deficits after induction with intranasal **α**-MSH (Trial 2a). Six months after induction, we observed neither reduction in DNs in the SNpc-VTA region (Figure 7a, [Supplementary Table II](https://drive.google.com/file/d/1Pbg4If9_5TSo4RrR-Jj8vJy7ks3_u0yI/view?usp=sharing)), nor reduction in striatal TH+ density (Figure 7b, [Supplementary Table III](https://drive.google.com/file/d/1sTRp6YIT1tcHaU2pZ4qxfmMopSgDupON/view?usp=sharing)), nor the presence of pSer129+ inclusions within the SNpc-VTA region in any of the mice (Figure 8, [Supplementary Table IV](https://drive.google.com/file/d/1Phd8QaCUz7as-j4QjDy-v7kdddi_PeCM/view?usp=sharing)). Rather, we observed accumulation of the p62 protein within the DNs in the SNpc-VTA region, confirming autophagic dysregulation (Table IV, [Supplementary Table V](https://drive.google.com/file/d/11lj6eTjg7CX5jRHZLBA1l5E7dvpqehjE/view?usp=sharing)). Moreover, we observed a 26% increase in the expression of TH protein in the striatum of mice exposed to elevated **α**-MSH as compared to striatum of control mice (Figures 9, [Supplementary Table III](https://drive.google.com/file/d/1sTRp6YIT1tcHaU2pZ4qxfmMopSgDupON/view?usp=sharing)).

**Figure 7:**
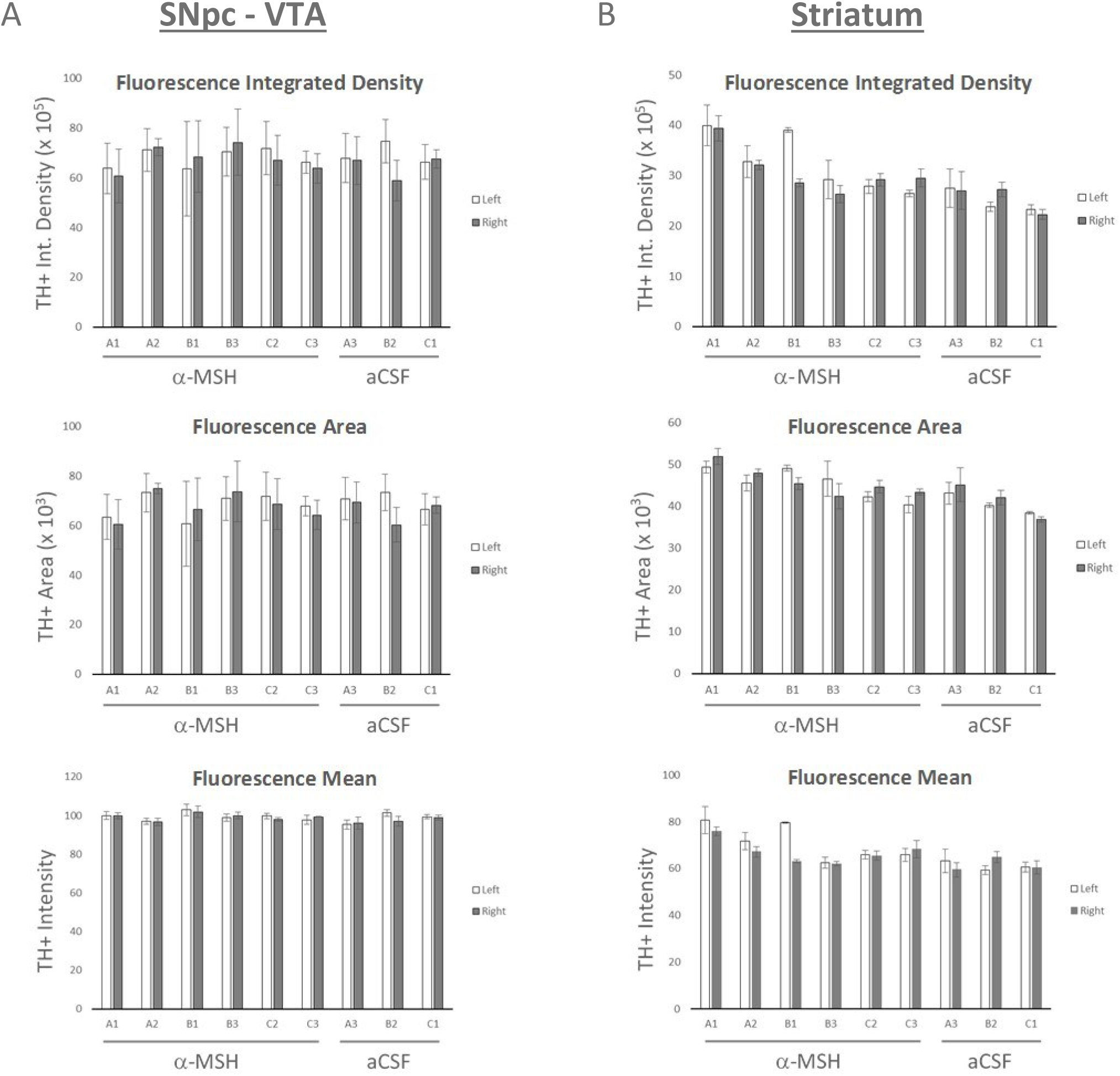
Substantia nigra pars compacta - VTA dopaminergic neurons and tyrosine hydroxylase positive striatal density remain intact in mice after intranasal *α*−MSH. Brains of mice (Trial 2a) were processed for histological assessment by immunofluorescent histochemistry. Fluorescence microscopy was used to obtain digital images of brain sections containing the (A) substantia nigra pars compacta – VTA region or (B) striatum stained with anti-tyrosine hydroxylase (TH) antibody. TH+ fluorescence was measured using the Image-J analysis software and expressed as integrated density, area and mean. Error bars are SEM of mean of triplicate sections. See Materials and Methods for experiment details and [Supplementary Table II](https://drive.google.com/file/d/1Pbg4If9_5TSo4RrR-Jj8vJy7ks3_u0yI/view?usp=sharing) and [Supplementary Table III](https://drive.google.com/file/d/1sTRp6YIT1tcHaU2pZ4qxfmMopSgDupON/view?usp=sharing), to view images and values used to calculate the means for (A) and (B), respectively.

**Figure 8:**
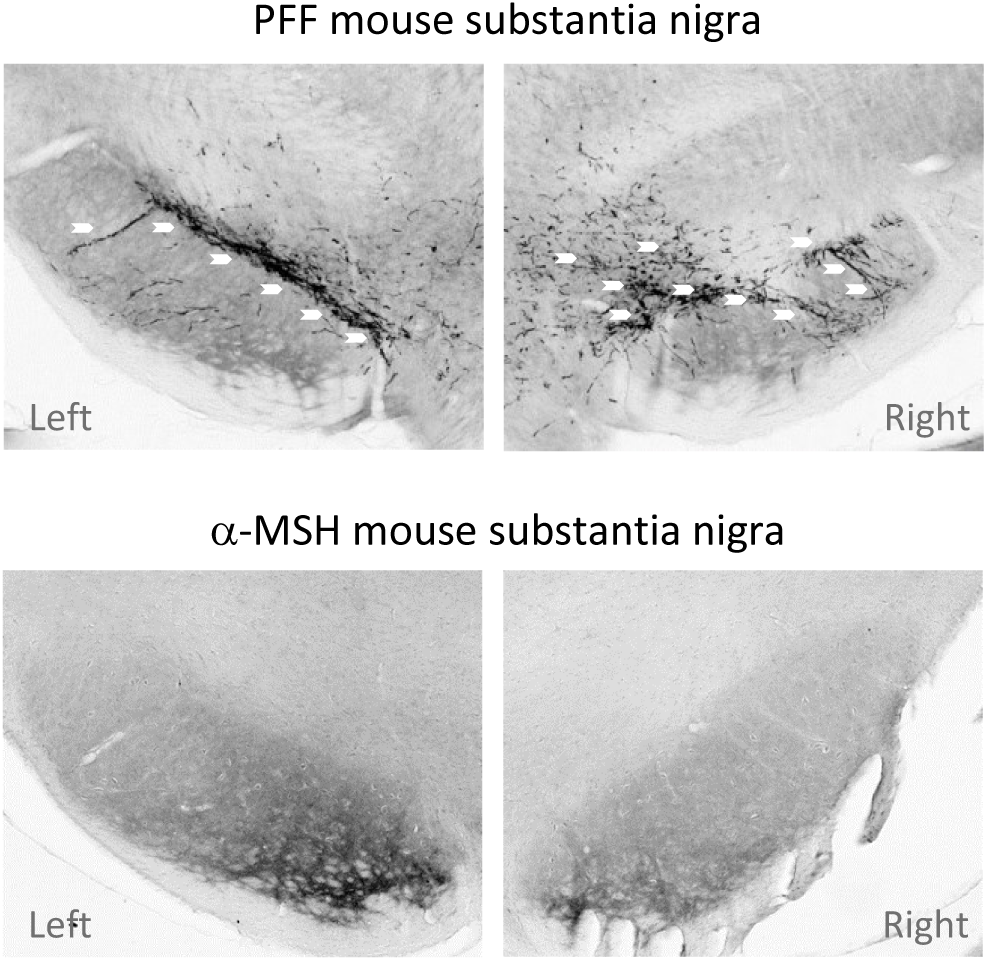
Alpha-synuclein inclusions are absent in the substantia nigra of mice administered *α*−MSH. Brains of mice (Trial 2a) were processed for histological assessment by immunohistochemisty and brain sections containing the substantia nigra were stained with anti-mouse alpha-synuclein (pSer129+) antibody. pSer129+ inclusions (arrows) were visible only in the brains of mice induced by injection with alpha-synuclein pre-formed fibrils (PFFs, top photos), and not in the brains of mice after 6 months of the one-month intranasal *α*−MSH administration (bottom photos, representative images). See Materials and Methods for experiment details and [Supplementary Table IV](https://drive.google.com/file/d/1Phd8QaCUz7as-j4QjDy-v7kdddi_PeCM/view?usp=sharing), to view additional images for each brain.

**Figure 9:**
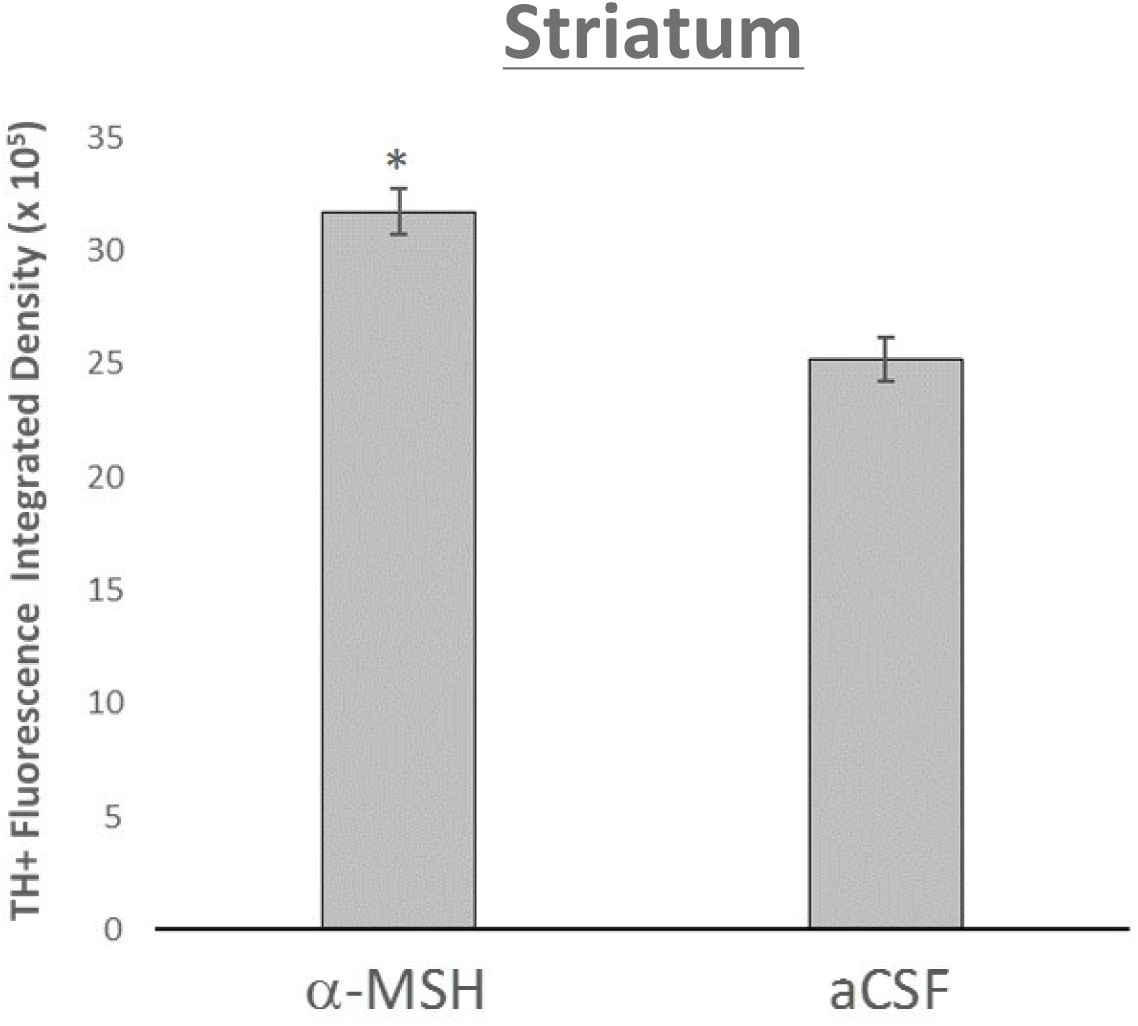
Increased striatal tyrosine hydroxylase expression in mice administered *α*−MSH. Tyrosine hydroxylase (TH)+ fluorescence integrated density measurements from left and right striatal sections (see also Figure 7b) were combined according to groups (*α*−MSH vs. aCSF) and the means compared. Error bars are SEM of mean of replicate measurements. A 26% increase in TH+ fluorescence integrated density was observed in striatal sections of mice administered *α*−MSH as compared to striatal sections from control mice (aCSF). *Welch’s T-test analysis of means and *p* value (two-tailed): *α*−MSH (n=36) vs. aCSF (n=18), *p* = 0.00002. A *p* value of less than 0.05 is considered significantly different. See Materials and Methods for experiment details and [Supplementary Table III](https://drive.google.com/file/d/1sTRp6YIT1tcHaU2pZ4qxfmMopSgDupON/view?usp=sharing), to view the images and values used to calculate the means.

**Table IV.**
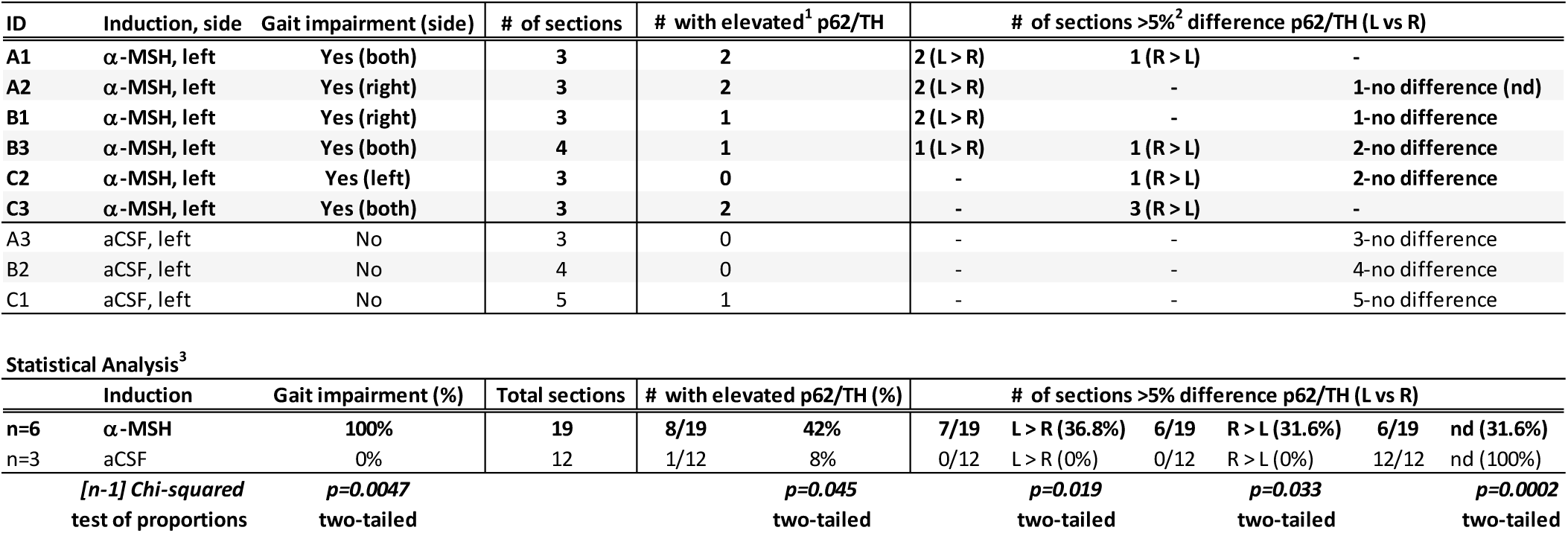
*α*−MSH induces autophagic dysfunction *in-vivo*: p62 accumulation in dopaminergic neurons of the substantia nigra pars compacta - VTA region. Brains of mice (Trial 2a) were processed for histological assessment by immunofluorescence histochemistry. Immunofluorescence microscopy was used to obtain digital images of brain sections containing the substantia nigra pars compacta (SNpc)-VTA regions. Images were analyzed using the Image-J software to measure the level of p62 protein within the dopaminergic neuronal cell population (TH+) (see [Supplementary Table V](https://drive.google.com/file/d/11lj6eTjg7CX5jRHZLBA1l5E7dvpqehjE/view?usp=sharing)). Consistent with the appearance of gait deficits in 100% of mice administered intranasal *α*−MSH, over 5Xs more sections had elevated p62 levels within the SNpc-VTA (TH+) region relative to the total number of sections as compared to the control mice (42% vs. 8%, respectively, *p* < 0.05), indicating autophagic dysfunction in dopaminergic neuronal cell population in these mice. Moreover, whereas the control brain sections had comparable p62/TH percentages on either side, 68.4% of sections from mice administered intranasal *α*−MSH had higher percentage of p62/TH expression on the left or right side (36.8% L > R, 31.6% R > L). This is consistent with the appearance of motor deficits that developed on either side, at equal frequency, even though induction by intranasal *α*−MSH was on the left nostril. **Footnotes:** ^1^ An unbiased histology analysis software was used to generate a border around the dopaminergic cells or tyrosine hydroxylase (TH+) positive cells within the SN-VTA region of the left and right brain sections stained with fluorescent labeled anti-TH (Cy3) and anti-p62 (Alexafluor 488) antibodies. Image-J analysis software was used to calculate the TH+ fluorescence area and p62+ fluorescence area within the marked border and the percentage of p62+ area was determined against TH+ area and considered elevated if p62/TH > 45%. See [Supplementary Table V](https://drive.google.com/file/d/11lj6eTjg7CX5jRHZLBA1l5E7dvpqehjE/view?usp=sharing) to view images, values obtained and calculations performed. ^2^ The p62/TH percentage was calculated for the left and right SN-VTA region for each section as described and sections that had a difference greater than 5% between the left or right SN-VTA region were counted and the side with the greater percentage of p62/TH reported. Sections that show a difference less than 5% between the left or right SN-VTA region were considered to have comparable p62/TH levels (no difference, nd). See [Supplementary Table V](https://drive.google.com/file/d/11lj6eTjg7CX5jRHZLBA1l5E7dvpqehjE/view?usp=sharing) to view images, values obtained and calculations performed. ^3^ [n-1] Chi-squared test of two proportions, expressed as a percentage, was used to determine if the two values are significantly different. A *p* value of less than 0.05 is considered significantly different.

## Discussion

We now propose a new mechanism that underlies the early events of dopaminergic dysfunction in PD. Our results suggest that exposure of MC1R-expressing DNs to elevated levels of microglial-derived **α**-MSH leads to impairment of cellular autophagy. Unabated, progressive cellular dysfunction gives way to abnormal accumulation of proteins, dopamine deficiency and motor deficits in PD.

The presence of activated microglial cells is a hallmark feature of PD pathology, which is detectable at the very early stage of the disease and is maintained through the duration of the disease [64–67]. During inflammation, activated microglial cells release **α**-MSH to maintain homeostasis [22, 68–71]. Consistent with persistent microglial cell activation in PD, **α**-MSH is elevated in the CSF of PD patients by at least two-fold above the baseline levels seen in control subjects [29, 30]. Similarly, in multiple system atrophy (MSA), a rare condition with symptoms and pathologies similar to PD, elevated CSF **α**-MSH correlates with increased microglial cell activation [29, 72]. During an acute traumatic brain injury (TBI), microglia are activated and the concentration of CSF **α**-MSH increases to modulate inflammation, peaking after 5-7 days [68]. With severe TBI, elevated levels of CSF **α**-MSH persist even longer by at least two-fold above that seen in healthy individuals [68]. TBI is implicated as a strong risk factor for the development of PD, with 56% increased risk after mild TBI to 83% increased risk after moderate to severe TBI [73, 74]. These observations implicate elevated CNS **α**-MSH levels in the development of PD and are consistent with our results, which show that prolonged CNS exposure to elevated **α**-MSH can give rise to dopaminergic dysfunction and lead to motor deficits in mice (Table I-IV, Figure 9).

Post-mortem analysis of the brains of individuals with incidental Lewy body disease (ILBD) showed alpha-synuclein pathology in the SN, despite having never shown signs of parkinsonism or dementia [75, 76]. Interestingly, microglia is elevated in both ILBD and PD SN compared to control subjects [75]. While microglial cell density is comparable between ILBD and PD, the increased presence of alpha-synuclein inclusions in PD correlated with an increased frequency of activated microglial cells [75, 77]. In ILBD, the microglial cell activation state is dominated by a pro-inflammatory M1 macrophage-like phenotype (TLR2+) [78]. In contrast to ILBD, this microglial cell subset is significantly lower in PD [75], which suggests that a pro-inflammatory milieu is neither the primary factor driving alpha-synuclein pathology nor the result of an increased presence of alpha-synuclein inclusions. Indeed, parkinsonian CSF had elevated anti-inflammatory compounds such as TGF-beta [79] and **α**-MSH [29, 30], which are both potent inducers of microglial cell activation toward the alternative M2-like phenotype [80, 81]. These observations suggests that the activation state of microglial cells predominant in PD reflects one that has had chronic exposure to elevated CSF **α**-MSH [29, 30].

While alpha-synuclein inclusions are suspected to drive degeneration in PD, deposits of Lewy bodies and neurites are not sufficient to produce parkinsonism, as seen in ILBD [12, 13, 75]. Despite adequate expression of alpha-synuclein pathology in both rodent and non-human primate models, motor deficits were absent [82]. A recent case report described an individual who was thoroughly examined and clinically diagnosed with PD but showed no alpha-synuclein inclusions in any brain region at post-mortem analysis. Rather, phosphorylated TDP-43 protein accumulation was abundant in the SN and in other areas of the brain [83]. These examples suggest that alpha-synuclein accumulation and aggregation may simply be an indicator of cellular dysfunction e.g., autophagic dysregulation, not the culprit that initiates or drives dopaminergic dysfunction in idiopathic PD.

Since the SN is densely populated by microglial cells [84, 85], DNs are vulnerable to exposure to elevated levels of **α**-MSH released by activated microglial cells during neuroinflammation. Patients with chronic neuroinflammation with a high probability to develop PD showed increased glucose consumption in the SN [86]. Similarly, we observed increased glucose consumption in MNT-1 cells after exposure to **α**-MSH (Figures 1, 2 and 3a). Our results show that exposure to **α**-MSH is apoptotic to cells under conditions of limited nutrients (Figures 3b and 3c), and the cells likely succumb to mitochondrial damage as a result of glucose deprivation and ATP depletion [87]. It is worth noting that in the advanced stages of PD, a decrease in glucose uptake is observed in the SN, reflecting neuronal degeneration and cell loss by apoptosis [6, 88]. Indeed, post-mortem analyses of PD SN reveal mitochondrial abnormalities, oxidative stress, high levels of reactive-oxygen species (ROS) and decreased glutathione levels. These conditions are found in tissues that experience glucose deprivation [87, 89–93].

While low glucose is normally a strong pro-autophagic stimulus (85), impairment of autophagy evoked by **α**-MSH was only surmountable by treatment with the MC1R antagonist and inverse agonist, ASIP (Figures 2, 3c and 4). This suggests that such “autophagic block” can be reversed. We observed progressive worsening of motor deficits in mice many weeks after ceasing intranasal **α**-MSH administration (Table II), suggesting that merely blocking **α**-MSH *in-vivo* may not be sufficient to reverse dopaminergic dysfunction active in these mice (Table III and IV). It may also require an inverse agonist like ASIP that, in addition to blocking **α**-MSH binding, transduces inhibitory signals at the MC1R receptor expressed on DNs [94–97].

Our **α**-MSH PD mouse model is an early-stage model lacking several features of late-stage PD pathology, namely dopaminergic neuronal cell loss, striatal dopamine axonal loss, and appearance of alpha-synuclein inclusions (Figures 7 and 8). Consistent with dopamine deficiency, autophagic dysregulation was observed in DNs in the SNpc-VTA region (Table IV), along with increased striatal TH expression (Figure 9). Increased TH expression suggests a compensatory early response to low dopamine levels [25, 98–100]. Resembling clinical symptoms of PD, **α**-MSH administered mice gradually developed movement deficits and rigidity (Table I-II), which were reversed with dopamine replacement and agonist treatment (Table III). The amount of time needed to observe end-stage pathology in this model might exceed the animal’s life span. However, in animals with a propensity to develop synucleinopathy, e.g., in transgenic mice expressing human alpha-synuclein (A53T), increased CNS exposure to **α**-MSH may shorten the time needed to observe end-stage pathology [101]. Nevertheless, the lack of late-stage features is not a weakness of the **α**-MSH PD mouse model, but a unique advantage that will be useful in testing and identifying drugs that act at the early stages of cellular dysfunction, thereby prevent diseased DNs from reaching a point when therapeutic intervention is no longer feasible.

A long standing knowledge in this field is that end-stage pathology hallmarks, such as significant dopaminergic cell loss and reduction in striatal TH density, are requisite for motor deficits in PD [102]. The mathematical models from which this is derived makes the critical assertion that significant dopaminergic cell loss (>30%) must be present to produce clinical symptoms of PD, namely, motor deficits. While our results do not agree with this paradigm, such bold assertion has had significant repercussions in the field of PD research. For example, disease modeling has been biased toward an end-stage disease phenotype that demands neuronal cell loss and reduction in striatal dopamine terminal volume. This is seen in the MPTP and 6-OHDA toxin animal models, which are currently used in PD drug research and development. Similarly, the alpha-synuclein PFF model has received much attention since the pathology it induces appears more consistent with the pathology observed in parkinsonian post-mortem tissue. Although it is still unclear what direct role, if any, alpha-synuclein inclusions have in PD pathophysiology, this feature is nevertheless considered cardinal in models of synucleinopathies and is perhaps a more desirable feature than motor symptoms!

End-stage disease models are used widely and have led to many new investigational drugs indicated to modify disease progression; however, these drugs are not effective in generating meaningful benefits for PD patients in clinical trials [103–105]. Employing end-stage PD models into drug design and selection has not translated successfully in clinical trials because the models likely do not represent the stage of pathogenesis most ideal for therapeutic intervention [106]. More early-stage disease model development, such as the work described here, are needed to usher in new therapies that will preserve neurons in the brain of patients.

Our results strongly suggest a central role for **α**-MSH at the onset of dopaminergic dysfunction in PD. Since **α**-MSH is implicated in more than one symptom of prodromal PD such as constipation [107, 108], restless leg syndrome [109, 110], sleep disorders [111–113] and excess sebum production “oily skin” [114, 115], **α**-MSH levels may serve as an early biomarker for disease onset and progression. Providing answers to this and other questions raised by our findings will help to produce a clearer understanding of PD pathophysiology.

## Acknowledgements

I would like to thank my wife, Linh Dela Cruz, who battles PD daily, for her love and support. I am thankful for the prayers and support of our families and friends. Thank you, Lord Jesus Christ, for your kindness to us all and for guiding us through this work. May all the glory be Yours alone. - JSDC

## Supplementary Materials

**Figures:**

[Supplementary Figure 1](https://drive.google.com/file/d/1s5mTF8NlumnD4xRiTB2gxRy6hcKeUfRP/view?usp=sharing)

[Supaplementary Figure 2](https://drive.google.com/file/d/1tFwqvDZiiPwewzBPok5ONSH2jMTKLrIa/view?usp=sharing)

**Tables:**

[Supplementary Table I](https://drive.google.com/file/d/1TuqhiNMlMDRwOQZe-Wx2A4lH15EBa1FG/view?usp=sharing)

[Supplementary Table II](https://drive.google.com/file/d/1Pbg4If9_5TSo4RrR-Jj8vJy7ks3_u0yI/view?usp=sharing)

[Supplementary Table III](https://drive.google.com/file/d/1sTRp6YIT1tcHaU2pZ4qxfmMopSgDupON/view?usp=sharing)

[Supplementary Table IV](https://drive.google.com/file/d/1Phd8QaCUz7as-j4QjDy-v7kdddi_PeCM/view?usp=sharing)

[Supplementary Table V](https://drive.google.com/file/d/11lj6eTjg7CX5jRHZLBA1l5E7dvpqehjE/view?usp=sharing)

**Videos:**

[Supplementary Video 1](https://drive.google.com/file/d/1SP0ngmMquihkBPMfnOwefMPA_CAFYOxL/view?usp=sharing)

[Supplementary Video 2](https://drive.google.com/file/d/1d9BS1Aw1jNj_Ynnrjox2Qf1KK0TltVsQ/view?usp=sharing)

[Supplementary Video 3](https://drive.google.com/file/d/161dcaztb__K7k5SqYJh8Eo4WQvd_FQUc/view?usp=sharing)

[Supplementary Video 4](https://drive.google.com/file/d/1Ghgz-Rxz4VTxrWextkhlmxqdyJ9B2yGx/view?usp=sharing)

[Supplementary Video 5](https://drive.google.com/file/d/1n7P1i6cCJgNymI5EXxtExPRullxHvap1/view?usp=sharing)

[Supplementary Video 6](https://drive.google.com/file/d/1gKyPdByOPbZBVwBpE7zttzteAmNzYBgS/view?usp=sharing)

[Supplementary Video 7](https://drive.google.com/file/d/1bZqG_mj3-5kAIFiYKrVE-NewvMlVMWLw/view?usp=sharing)

[Supplementary Video 8](https://drive.google.com/file/d/1e1tam5SbYJNDNMa6vWGgxFvQxliPcZ79/view?usp=sharing)

[Supplementary Video 9](https://drive.google.com/file/d/1Ihodmi4X-_RKz8vzqtb3JGpSI6yn3y61/view?usp=sharing)

[Supplementary Video 10](https://drive.google.com/file/d/1dc7kGG-1JTrNx3HOOUC4gghOmHs_NDwX/view?usp=sharing)

[Supplementary Video 11](https://drive.google.com/file/d/1mzfehtT0xv-TSScSPOSVQ59NzV5in7H1/view?usp=sharing)

[Supplementary Materials List](https://drive.google.com/file/d/1J_M3L_jcLMSrtDKkhd0ZUivkzsvDPx9e/view?usp=sharing)

